# Crosstalk between H2A variant-specific modifications impacts vital cell functions

**DOI:** 10.1101/2021.01.14.426637

**Authors:** Anna Schmücker, Bingkun Lei, Zdravko J. Lorković, Matías Capella, Sigurd Braun, Pierre Bourguet, Olivier Mathieu, Karl Mechtler, Frédéric Berger

## Abstract

Histone variants are distinguished by specific substitutions and motifs that might be subject to post-translational modifications (PTMs). Compared with the high conservation of H3 variants, the N- and C-terminal tails of H2A variants are more divergent and are potential substrates for a more complex array of PTMs, which have remained largely unexplored. We used mass spectrometry to inventory the PTMs of the two heterochromatin-enriched variants H2A.W.6 and H2A.W.7 of *Arabidopsis*, which harbor the C-terminal motif KSPK. This motif is also found in macroH2A variants in animals and confers specific properties to the nucleosome. We showed that H2A.W.6 is phosphorylated by the cell cycle-dependent kinase CDKA specifically at KSPK. In contrast, this modification is absent on H2A.W.7, which also harbors the SQ motif associated with the variant H2A.X. Phosphorylation of the SQ motif is critical for the DNA damage response but is suppressed in H2A.W.7 by phosphorylation of KSPK. To identify factors involved in this suppression mechanism, we performed a synthetic screen in fission yeast expressing a mimic of the *Arabidopsis* H2A.W.7. Among those factors was the BRCT-domain protein Mdb1. We showed that phosphorylation of KSPK prevents binding of the BRCT-domain protein Mdb1 to phosphorylated SQ and as a result hampers response to DNA damage. Hence, cross-talks between motif-specific PTMs interfere with the vital functions of H2A variants. Such interference could be responsible for the mutual exclusion of specific motifs between distinct H2A variants. We conclude that sequence innovations in H2A variants have potentiated the acquisition of many specific PTMs with still unknown functions. These add a layer of complexity to the nucleosome properties and their impact in chromatin regulation.

## Introduction

Histones represent the major protein component of chromatin. Histone variants evolved in all core histone families and acquired comparable properties in a convergent manner [1–4]. These variants play major roles in cell fate decisions, development, and disease [5–7]. Most multicellular eukaryotes contain three types of H2A variants: H2A, H2A.Z and H2A.X. The variant H2A.X is defined by the motif SQ[E/D]Φ present within the C-terminal tail (where Φ stands for a hydrophobic amino acid) that is present only at the C-terminal tail of H2A.X. The serine residue of this motif is phosphorylated during the early phase of the DNA damage response (DDR) [7–11]. In animals, serine 139 (S139) phosphorylation at the SQ motif of H2A.X (γH2A.X) is sufficient to recruit the mediator of DNA damage checkpoint protein 1 (MDC1) [12]. In contrast, MDC1 binding is abolished if tyrosine 142 (Y142) of γH2A.X is phosphorylated. Thus, the succession of these two phosphorylation steps dictates the order of events at sites of DNA damage [13–15]. In plants and yeast, serine phosphorylation of SQ[E/D]Φ is also essential for DDR although Y142 of H2A.X is not conserved [16, 17]. At the initiation of DDR in fission yeast, γH2A.X is recognized by the BRAC1 C-terminal (BRCT) domains of Crb2 [18] and Mdb1, the ortholog of human MDC1 [19]. These events are genetically redundant and initiate formation of radiation-induced nuclear foci at the site of DNA damage, which act as template for recruiting DNA repair machinery. BRCT domain-containing proteins also exist in plants, yet their functions remain unknown.

In *Arabidopsis*, in addition to H2A.X, DDR also relies on the plant specific variant H2A.W.7 that harbors an SQ motif at the C-terminal tail [9, 16, 20]. The family of H2A.W variants evolved in land plants, where they exclusively occupy constitutive heterochromatin, and consists of three isoforms in *Arabidopsis* (H2A.W.7, W.6 and W.12) [2, 16, 21]. H2A.W confers distinct properties to the nucleosome through differences in its primary amino acid sequence in the L1 loop, the docking domain, and the extended C-terminal tail [22]. The main feature defining H2A.W variants is the C-terminal KSPK motif [21, 23]. Due to the expected location of the H2A.W C-terminus at the nucleosome dyad (entry/exit site of the DNA into the nucleosome), the KSPK motif is placed in a functionally significant area [24] where it interacts with the linker DNA [25]. This motif is a member of the S/T-P-X-K/R motifs (where X represents any amino acid) present in macroH2A in metazoans and linker histones H1 among other proteins [26–32]. Both H2A.W and macroH2A are required for heterochromatin organization, suggesting a potential convergence in the function of these histone variants [2]. Incorporation of these variants confers specific biophysical properties to the nucleosomes and chromatin [1, 26, 33, 34].

Distinct C-terminal motifs present in specific classes of H2A variants do not usually co-occur. The KSPK is present exclusively in plant H2A.W and mammalian macroH2A, while the SQ[E/D]Φ motif is present primarily on H2A.X variants in eukaryotes [5, 9, 33, 35]. There are notable exceptions to this rule in animals and plants. *Drosophila* H2A.V combines the SQ[E/D]Φ motif with properties of H2A.Z [36]. Several species of seed-bearing plants possess a subtype of H2A.W that also harbors a SQ[E/D]Φ motif [16]. In *Arabidopsis*, H2A.X is largely excluded from constitutive heterochromatin, which is occupied by the variant H2A.W.7 that carries both the SQ[E/D]Φ and KSPK motifs. H2A.X and H2A.W.7 are essential to mediate the response to DNA damage in *Arabidopsis*, but variants similar to H2A.W.7 are present only in a restricted number of flowering plant species [16]. What led to the mutual exclusion of C-terminal motifs during H2A variant evolution has remained unclear, but one reason could be incompatibility between post-translational modifications (PTMs) on variant specific motifs.

Different types and combinations of PTMs of core histones coordinate the recruitment of proteins that dictate chromatin configuration and consequently regulate genome integrity and genome expression [5, 7, 15, 37–39]. Non-centromeric H3 variants share strong sequence homology and, with a few exceptions [40–42], are subjected to the same repertoire of modifications. In contrast, sequence homology between H2A variants, particularly at their N- and C-terminal tails, is much less pronounced [23, 33]. This provides opportunities for deposition of distinct patterns of modifications for each type of H2A variant [7, 43].

Here, we provide an inventory of PTMs of H2A.W variants in *Arabidopsis*, including some that are specific of subtypes of H2A.W variants. H2A.W.6 is phosphorylated at the serine residue of the KSPK motif, and we provide evidence that the modification is deposited by cyclin dependent kinases (CDKs). In contrast, H2A.W.7 is only phosphorylated on the SQ motif. Notably, H2A.W.7 carrying a phosphomimetic KDPK motif shows impaired DDR. Through a synthetic approach in fission yeast, we show that phosphorylation of the KSPK motif prevents Mdb1 binding to the phosphorylated SQ motif and proper DDR, suggesting that the absence of KSPK phosphorylation in H2A.W.7 is essential for the SQ motif to mediate DDR in *Arabidopsis*. Hence, PTMs of C-terminal motifs of the H2A.W.7 variant interfere with each other and, during DDR this seems to be resolved by suppression of PTMs on the KSPK motif. Yet, this type of variant remains rarely present in eukaryotes. We propose that incompatibility between the PTMs carried by H2A variant specific motifs provide a possible explanation for the mutual exclusion of C-terminal motifs between H2A variants.

## Results

### H2A.W.6 and H2A.W.7 display distinct patterns of modifications at their C-terminal tails

We immunopurified mononucleosomes containing H2A.W.6 or H2A.W.7 from wild type (WT) leaves of *Arabidopsis* (Fig 1A and 1B) and performed qualitative MS analysis to identify PTMs associated with each isoform. In both isoforms, N- and C-terminal tails were modified at several lysine residues by acetylation and/or methylation (Fig 1C). A more complex set of modifications was found on the C-terminal tail of H2A.W.6, with prevalence of lysine acetylation and serine phosphorylation (Fig 1C). Although H2A.W.6 and H2A.W.7 exist primarily as homotypic (two copies of either H2A.W.6 or H2A.W.7), heterotypic (one copy of each H2A.W.6 and H2A.W.7) nucleosomes can be identified [16, 22] (Fig 1B). Importantly, H2A.W.6 and H2A.W.7 precipitated in respective and reciprocal immunoprecipitations contained similar sets of modifications (S1 Fig), suggesting that each variant isoform acquires distinct modification patterns independently of the nucleosome composition. Some of these modifications were specific to each isoform, in part due to the absence of conservation of the residues targeted by these modifications. Three lysine residues at the N-terminal tails of H2A.W.6 and H2A.W.7 present in a highly conserved sequence context (KSVSKS<u>M</u>KAG vs. KSVSKS<u>V</u>KAG) showed similar PTMs in both variants. In contrast, other lysine residues at the N-termini in a less conserved context displayed isoform-specific modifications (Fig 1C). On both variants, acetylation was detected on lysine residue 128 and 144, which are embedded in the same sequence context (Fig 1C). Seedlings, leaves and flowers showed similar patterns of lysine modifications at the C-terminal tail, but they differed in their range and abundance. The three serine residues (S129, S141, and S145) at the C-terminal tail were phosphorylated on H2A.W.6 in leaves, as previously reported [44]. Based on spectral counting, the most abundant phosphorylation was S145, followed by S141 and S129; the last one being very rare. Phosphorylation of S141 and S145 were also detected on H2A.W.6 from seedlings, where a single spectrum also detected S129 phosphorylation of H2A.W.7. In flowers, only S145 was detected on H2A.W.6 (S1 Fig). Overall, the repertoire of PTMs detected on H2A.W.6 and H2A.W.7 differed markedly (Fig 1C). As S145 of the conserved KSPK motif is part of the functionally relevant C-terminal tail that protects the linker DNA [22, 23], we focused our further analysis on S145 phosphorylation. We obtained an antibody that specifically binds phosphorylated S145 in both H2A.W.6 and H2A.W.7 *in vitro* (S2 Fig). Yet, consistent with the MS data, *in vivo* KSPK phosphorylation was only detected on H2A.W.6 (Fig 1C and 1D) and we therefore named this antibody H2A.W.6p. Our data suggest that phosphorylation of the KSPK motif is deposited on H2A.W.6 but not on H2A.W.7, which can be phosphorylated on its SQ motif in response to DNA damage.

**Fig 1.**
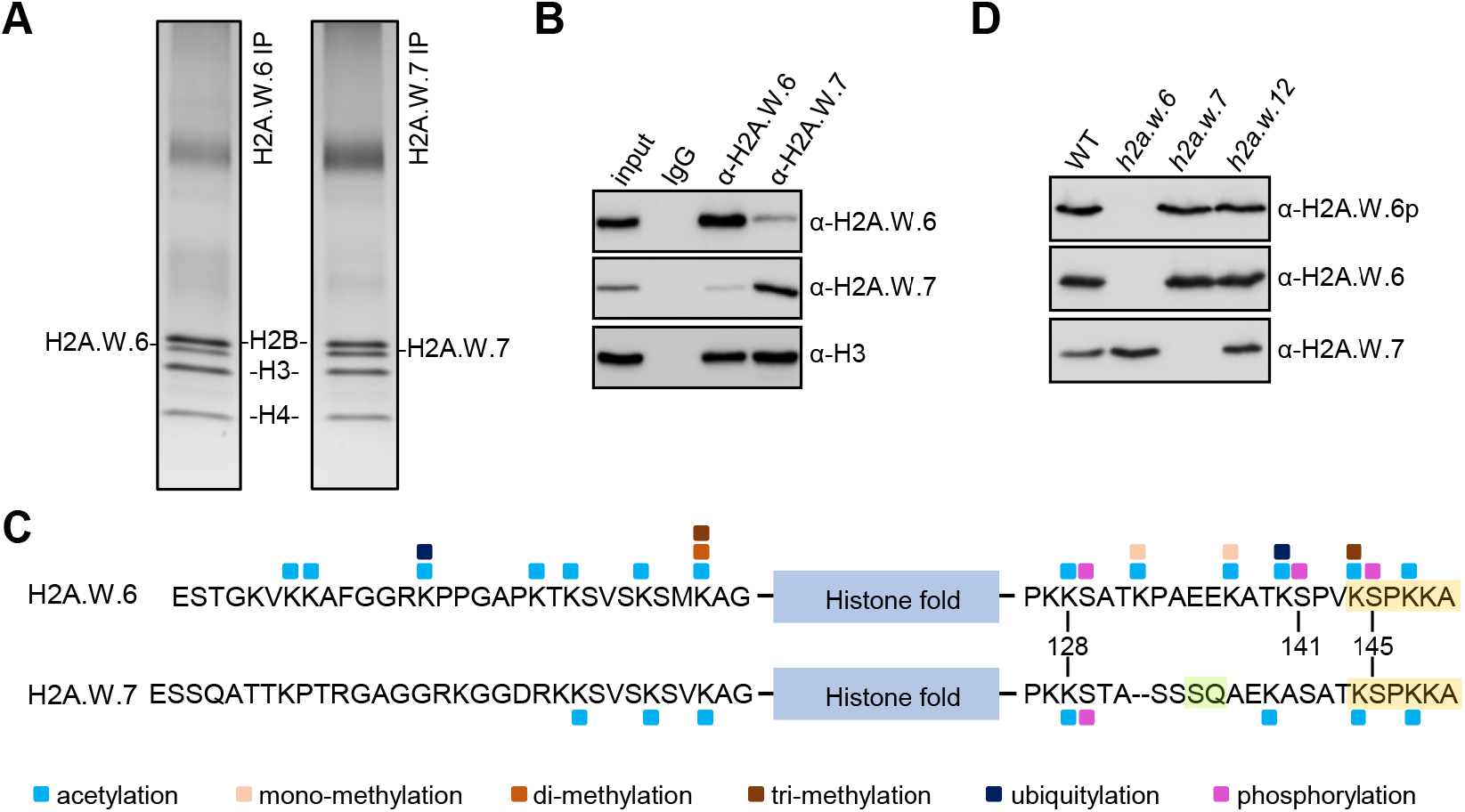
H2A.W.6 and H2A.W.7 carry distinct PTMs. (A) Silver stained gels of H2A.W.6 and H2A.W.7 mononucleosomes immunoprecipitated from MNase digested nuclear extracts from leaves. Bands corresponding to H2A.W.6 and H2A.W.7 were excised and analyzed by mass spectrometry (MS). (B) Western blot analysis of immunoprecipitates obtained with H2A.W.6 and H2A.W.7 specific antibodies from MNase digested WT nuclei. Note that H2A.W.6 and H2A.W.7 nucleosomes contain small amounts of H2A.W.7 and H2A.W.6, respectively, as previously reported [16, 22]. (C) Summary of all PTMs detected on H2A.W.6 and H2A.W.7. Amino acid sequence of N- and C-terminal tails of H2A.W.6 and H2A.W.7 are indicated with the conserved H2A.W KSPK motif and H2A.W.7 SQ motif highlighted in orange and green, respectively. The blue box indicates the histone fold domain. Post-translational modifications detected by MS are color-coded as indicated at the bottom. (D) Western blot analysis of nuclear extract from twelve days old WT, *hta6, hta7*, and *hta12* mutant seedlings with antibodies against H2A.W.6p, H2A.W.6, and H2A.W.7.

### The KSPK motif of H2A.W.7 is not phosphorylated upon DNA damage induction

We investigated the modification of KSPK and SQ motifs in H2A.W.6, H2A.W.7, and H2A.X after inducing DNA damage with bleomycin treatment. In WT, bleomycin treatment induced phosphorylation of H2A.X and H2A.W.7 at the SQ motifs (γH2A.X and γH2A.W.7), as previously reported [16] (Fig 2A). To determine whether bleomycin treatment induced phosphorylation of KSPK on H2A.W.7, we applied two-hour treatment with bleomycin to *hta6* mutant seedlings, which are deprived of H2A.W.6 and only expressed H2A.W.7 (Fig 2A). Under these conditions, KSPK phosphorylation was not detected, suggesting that DDR does not induce phosphorylation of KSPK on H2A.W.7. In conclusion, DNA damage triggers phosphorylation of the SQ motif of H2A.X and H2A.W.7, but not of the KSPK motif of H2A.W.7.

**Fig 2.**
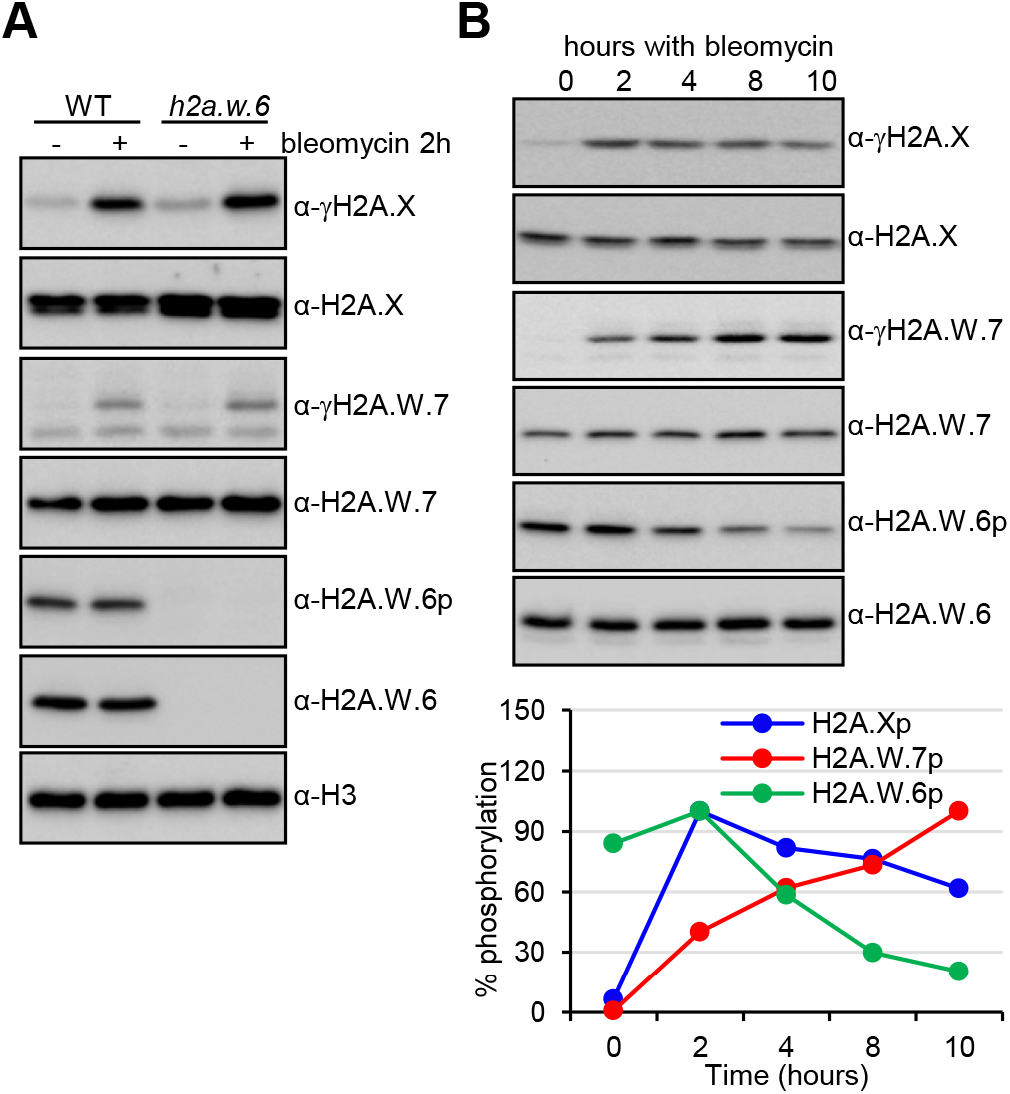
Phosphorylation of the H2A.W KSPK motif is not induced by DNA damage. (A) Phosphorylation of the KSPK motif on H2A.W.7 cannot be triggered by DNA damage. WT and *hta6* mutant seedlings were either mock or bleomycin (20 µg/ml) treated for two hours and nuclear extracts were analyzed by western blotting with indicated antibodies. (B) *Arabidopsis* WT seedlings were treated with 20 µg/ml of bleomycin for the indicated time periods and protein extracts were analyzed with the indicated antibodies. Levels of γH2A.X, γH2A.W.7, and H2A.W.6p (bottom panel) were quantified by normalization to the total level of the respective variant at each time point.

### H2A.W.6 phosphorylation is mediated by CDKA and is cell cycle dependent

We observed that prolonged exposure to bleomycin caused a marked decrease of KSPK phosphorylation of H2A.W.6 (Fig 2B). As DNA damage causes cell cycle arrest [45, 46], we hypothesized that KSPK phosphorylation could be associated with cell cycle progression. Because our *Arabidopsis* cell suspension could not be synchronized, we used tobacco BY-2 cell suspension for cell cycle synchronization to address this question. In tobacco, six of the seven H2A.W isoforms contain the KSPK motif at the C-terminus, of which phosphorylation was detected by the H2A.W.6p antibody (S3A and S3B Fig). We analyzed H2A.W phosphorylation status in synchronized tobacco BY-2 cells over a twelve-hour time course after release from an aphidicolin-induced cell cycle block (S3C and S3D Fig). While H2A.W.6 levels were comparatively stable throughout the cell cycle, phosphorylation of KSPK remained stable during S and G2 but decreased during G1 phase (Fig 3A and S3E Fig). We tested whether levels of H2A.W.6p also fluctuated in dividing cells of *Arabidopsis* root tips where pairs of small flat cells in G1 are easily distinguished from larger cells in G2 phase (Fig 3B). Cells in S phase were marked by a pulse of EdU incorporation. Consistent with results obtained with synchronized BY-2 cells, *Arabidopsis* root tip nuclei in S, M, and G2 phases showed high levels of H2A.W.6 phosphorylation, whereas this mark was not detected in G1 phase nuclei (Fig 3B). This fluctuation was not the result of changes in total levels of H2A.W.6, which remained relatively uniform throughout the cell cycle (S4A Fig). These results showed that phosphorylation of H2A.W.6 is dependent on cell cycle progression.

**Fig 3.**
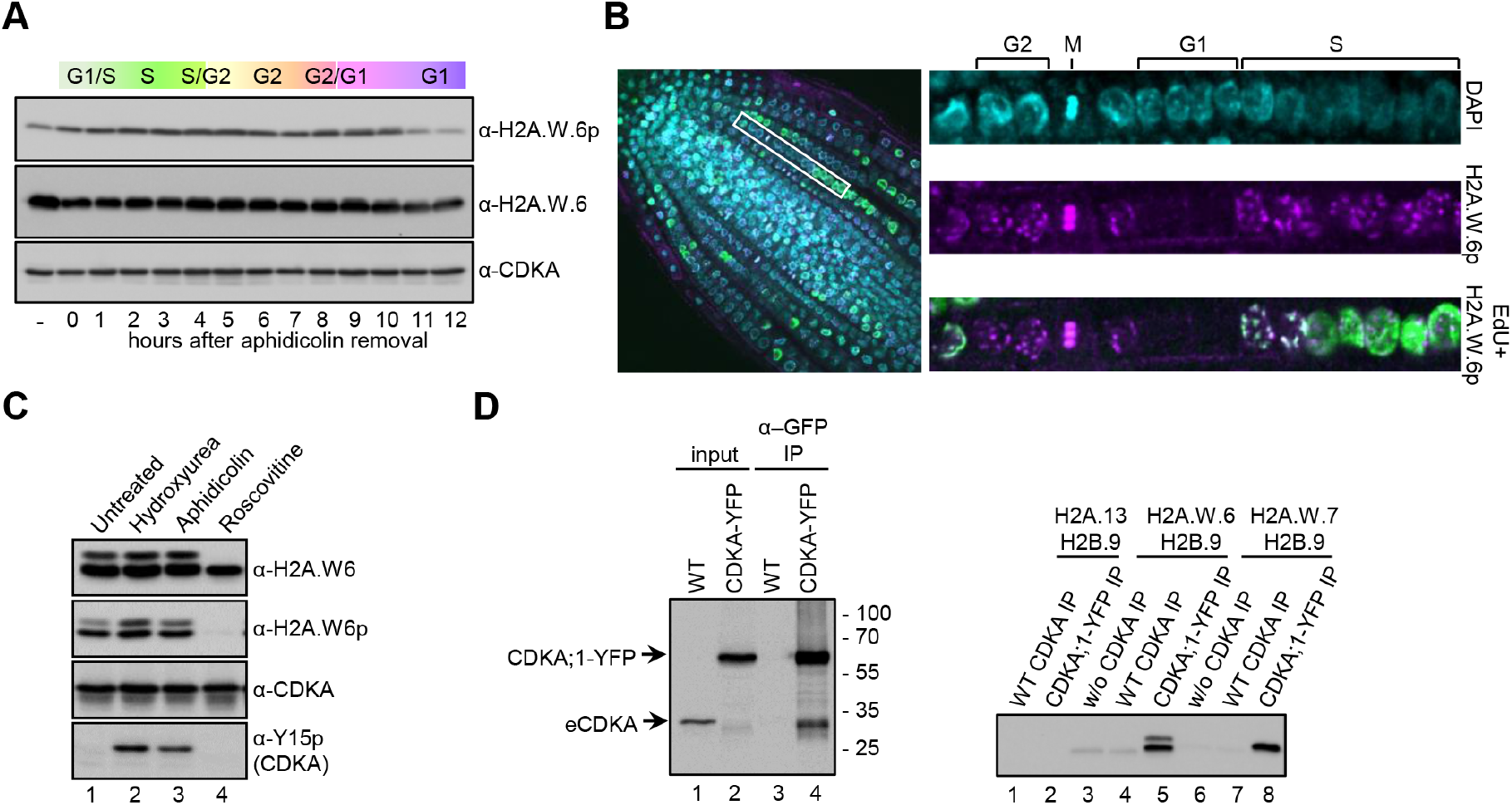
H2A.W.6 is phosphorylated in a cell cycle dependent manner by cyclin dependent protein kinases (CDK). (A) Phosphorylation of the KSPK motif is cell cycle dependent in tobacco BY-2 cell suspension culture. BY-2 cells were synchronized with 20 µg/ml aphidicolin for 24 hours. Protein extracts from samples taken in one-hour intervals after release of the block were analyzed by western blotting with the indicated antibodies. (B) Confocal images of WT root tips immunostained with H2A.W.6p antibody (magenta) after EdU (green) incorporation. Enlarged images of a row of cells in different cell cycle stages (indicated on the top) are shown on the right. (C) Protein extracts from *Arabidopsis* cell suspension treated with cell cycle inhibitors were analyzed by western blotting with the indicated antibodies. Inhibitory phosphorylation of CDK at tyrosine 15 (Y15) upon hydroxyurea and aphidicolin treatment (bottom panel, lanes 2 and 3) demonstrate the specificity and robustness of the assay. By contrast, roscovitine that inhibits the CDK activity directly by binding to the ATP binding pocket does not induce Y15 phosphorylation. (D) *Arabidopsis* CDKA;1-YFP was immunoprecipitated from whole cell extracts and detected with an anti-CDK antibody (left panel, lanes 2 and 4) and used in an *in vitro* kinase assay (right panel) with recombinant histone H2A-H2B dimers as indicated. Phosphorylation of the KSPKK motif was detected by western blotting with H2A.W.6p antibody. Note that GFP beads do not precipitate wild-type CDKA;1 (left panel, lane 3) and consequently H2A.W.6p was not detected in kinase assays from these IPs (right panel, lanes 4 and 7).

We next tested the impact of cell cycle inhibition on H2A.W.6p levels. S phase arrest by treatment with hydroxyurea or aphidicolin did not affect H2A.W.6 phosphorylation in *Arabidopsis* cell suspension cultures (Fig 3C). In contrast, when we inhibited cyclin-dependent kinases (CDKs) with roscovitine [47], H2A.W.6p was almost undetectable (Fig 3C). The specificity and concentration dependence of H2A.W.6p inhibition by roscovitine (Fig 3C, S4B Fig) prompted us to examine the *Arabidopsis* cyclin-dependent kinase CDKA;1, which is predominantly active at the G1 to S-phase transition [48–50]. We immunopurified CDKA;1-YFP from *Arabidopsis* seedlings and performed an *in vitro* kinase assay with recombinant H2A.W.6-H2B, H2A.W.7-H2B, and H2A-H2B dimers (Fig 3D). We observed strong phosphorylation of the H2A.W.6-H2B and H2A.W.7-H2B dimers but no signal in the H2A-H2B control (Fig 3D), demonstrating that CDKA;1 phosphorylates H2A.W.6 and H2A.W.7 *in vitro*. In plants with reduced CDKA;1 kinase activity [51, 52], we detected very low levels of H2A.W.6 phosphorylation (S4C Fig), further supporting that CDKA;1 is the main kinase responsible for H2A.W.6 modification. In root tips of plants with reduced CDKA;1 kinase activity, only late G2/M-phase nuclei showed KSPK phosphorylation (S4D Fig). This suggested that another CDK phosphorylates the KSPK motif in the absence of CDKA;1, which might be CDKB;1, as it is active at the G2/M transition [49, 50, 53].

In conclusion, H2AW.6 is phosphorylated by cyclin-dependent kinases during the S, G2, and M phases and the modification is likely removed during the G1 phase of the cell cycle.

### Cross talk between phosphorylation at the KSPK and SQ motifs in H2A.W.7

We demonstrated that CDKA;1 phosphorylates H2A.W.7 at the KSPK motif *in vitro* (Fig 3D); however, this modification was not detected *in planta* (Fig 1C and 1D, S2 Fig). We thus tested whether DNA damage induced phosphorylation of the SQ motif interferes with KSPK phosphorylation in H2A.W.7. This was not the case in *hta7* plants expressing phosphomimetic (SQ to DQ) mutant forms of H2A.W.7 [16]. Additionally, KSPK phosphorylation was also not observed when SQ phosphorylation was prevented in *hta7* plants expressing non-phosphorylatable (SQ to AQ) (S5A Fig). Hence, phosphorylation of the H2A.W.7 SQ motif does not appear to interfere with KSPK phosphorylation.

To test the crosstalk between SQ and KSPK phosphorylation *in planta*, we attempted to complement the *hta7* mutant by expressing WT and mutant forms of H2A.W.7 (Fig 4A). Expression of a mutant form of H2A.W.7 combining the WT SQ motif with a KDPK motif that mimics phosphorylation (SQ-DP) did not rescue sensitivity of *hta7* mutant plants towards DNA damage (Fig 4B). By contrast, mutation of the KSPK motif into non-phosphorylable KAPK (SQ-AP) rescued DDR (Fig 4B). Notably, mutations of the KSPK motif into either KAPK or KDPK did not affect SQ motif phosphorylation (Fig 4C). Thus, the presence of a negative charge at the KSPK motif does not interfere with SQ motif phosphorylation but causes DNA damage sensitivity. These results suggested that serine phosphorylation of KSPK must be prevented to execute DDR.

**Fig 4.**
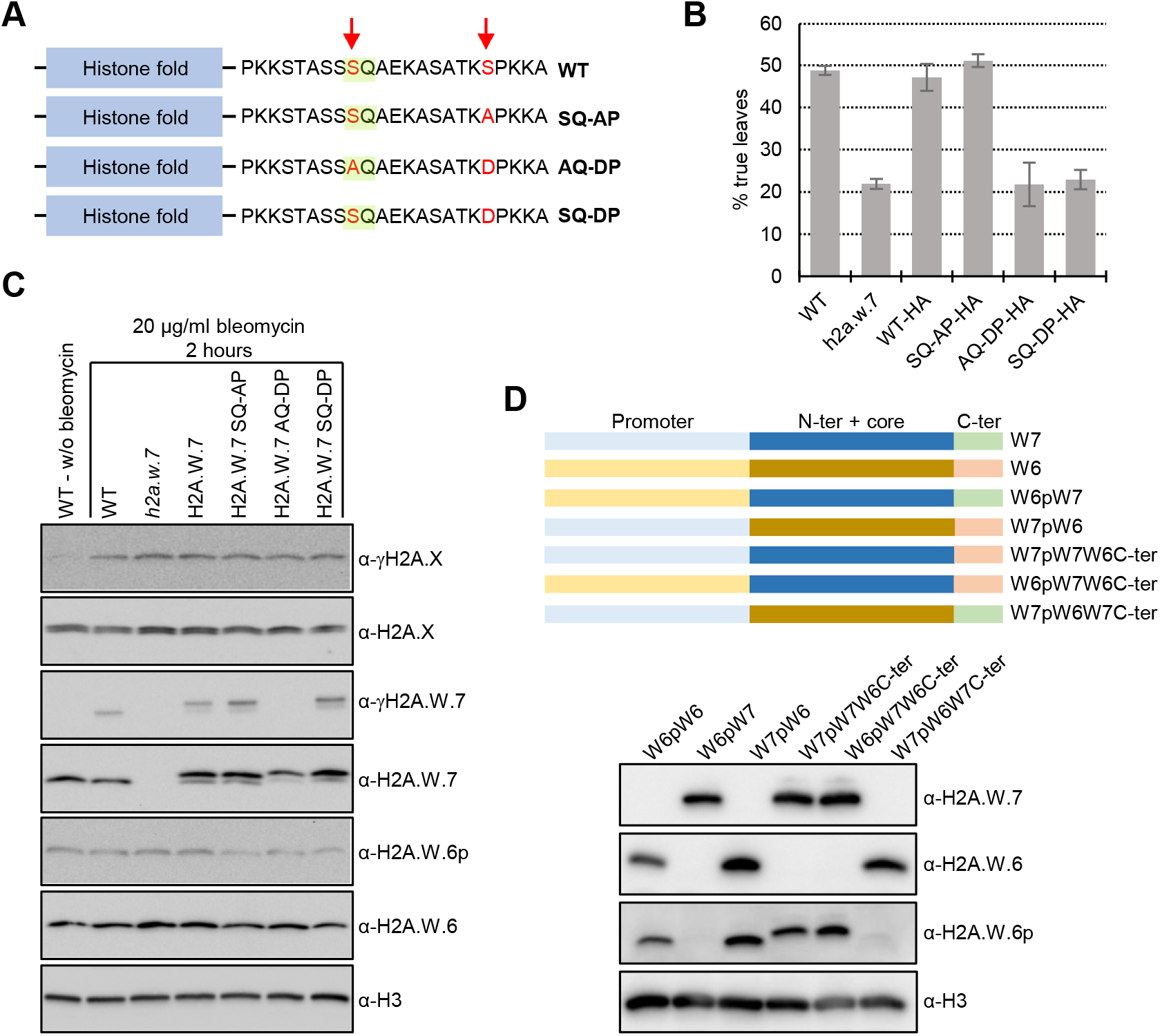
Phosphomimic of KSPK phosphorylation results in DNA damage sensitivity *in planta*. (A) Schematic diagrams of *Arabidopsis* H2A.W.7 variants containing mutated serin residues (in red) in the SQ (highlighted in green) and KSPK motifs. (B) Phosphomimic of the KSPK motif confers DNA damage sensitivity. Seeds from *h2a*.*w*.*7* mutants expressing HA-tagged WT H2A.W.7 or H2A.W.7 with mutations in the SQ and KSPK motifs were germinated in the presence of 50 mg/mL of zeocin and scored for true leaf development twelve days after germination. Data are represented as means ±SD of three independent experiments with n>400 seedlings. (C) Phosphomimic of the KSPK motif does not prevent SQ motif phosphorylation of H2A.W.7. Seedlings grown for 10 days on MS plates were treated for two hours with 20 µg/ml of bleomycin and nuclear extracts were analyzed by western blotting with the indicated antibodies. (D) Primary sequence of the H2A.W C-terminal tail determines phosphorylation outcome of the KSPK motif. Schemes of promoter, histone core domain and C-terminal tail swaps between H2A.W.6 and H2A.W.7 (top panel). Western blot analysis of expression and KSPK motif phosphorylation of nuclear extracts from plants expressing indicated H2A.W swap versions in mutant plants deprived from H2A.W (bottom panel).

What are the factors preventing phosphorylation of KSPK in H2A.W.7? We hypothesized that the pattern of expression of H2A.W.6 and H2A.W.7 are different leading to expose these two variants to contrasting activities that deposit and maintain KSPK phosphorylation. Contrary to our hypothesis, we observed that KSPK phosphorylation in H2A.W.7 was still absent even when expressed under the control of the promoter of H2A.W.6 in the *hta7* mutant (Fig 4D). Residues surrounding the KSPK motifs are distinct between H2A.W.6 and H2A.W.7. Consistently, we observed phosphorylation of KSPK of the C-terminal tail of H2A.W.6 when fused to the core of H2A.W.7 (Fig 4D). This suggested that the primary sequence of the C-tail of H2A.W.6, but not of H2A.W.7, is prone to be phosphorylated *in planta*.

### Cross talk between phosphorylation at the KSPK is mediated by proteins containing a BRCT domain that binds to phosphorylated SQ

To further investigate a potential crosstalk between phosphorylation of the H2A.W.7 SQ and KSPK motifs, we took a synthetic approach using the fission yeast *Schizosaccharomyces pombe*. Fission yeast possesses a relatively reduced repertoire of histone H2A variants, consisting of two H2A.X variants (*Sp*H2A.α and *Sp*H2A.β) and H2A.Z [54, 55]. We modified both genes encoding *Sp*H2A by inserting the C-terminal tail of the *Arabidopsis* H2A.W.6 to the C-terminus of *Sp*H2A.α and *Sp*H2A.β that contains an SQ motif, and obtained the chimeric histone *Sp*H2A.W^*At*^ that possesses the motifs present in H2A.W.7 [25] (Fig 5A, S5B Fig). In dividing cells, *Sp*H2A.W^*At*^ localized to the nucleus and was incorporated in chromatin (Fig 5B, S5C Fig). We detected phosphorylation of *Sp*H2A.W^*At*^ at the KSPK motif but not in a control strain that expressed *Sp*H2A.W^*At*^ where KSPK was substituted by four alanine residues (*Sp*H2A.W4A^*At*^) (Fig 5A and 5C). Interestingly, phosphorylation at the KSPK and SQ motifs were both detected on *Sp*H2A.W^*At*^ by mass spectrometry (Fig 5E, S5D Fig), demonstrating that *Sp*H2A.W^*At*^ phosphorylation did not prevent SQ phosphorylation. While the yeast strain expressing *Sp*H2A.W^*At*^ lacks H2A.X, we predicted that SQ phosphorylation of *Sp*H2A.W^*At*^ would be sufficient to respond to DNA damage. However, expression of *Sp*H2A.W^*At*^ caused hyper-sensitivity to DNA damage (Fig 5D). This suggested that either extension of the C-terminal tail or modification of the KSPK motif interfered with the DDR normally mediated by phosphorylation of the SQ motif. To test these two possibilities, we examined DNA damage sensitivity of strains expressing either *Sp*H2A.W^*At*^, the KSPQ mutant *Sp*H2A.W4A^*At*^, *Sp*H2A with a repeated wild type C-terminal tail (*Sp*H2ACT; contains two SQ motifs) or with alanine substitution of the second SQ motif (*Sp*H2ACT-AA) (Fig 5A). Phosphorylation of the SQ motifs was detected in all strains and none of these modifications resulted in sensitivity to DNA damage treatment, except for *Sp*H2A.W^*At*^ cells (Fig 5D, S5D Fig). We thus concluded that the mere extension of the C-terminal tail is not responsible for the increased sensitivity to DNA damage in the strain expressing *Sp*H2A.W^*At*^.

**Fig 5.**
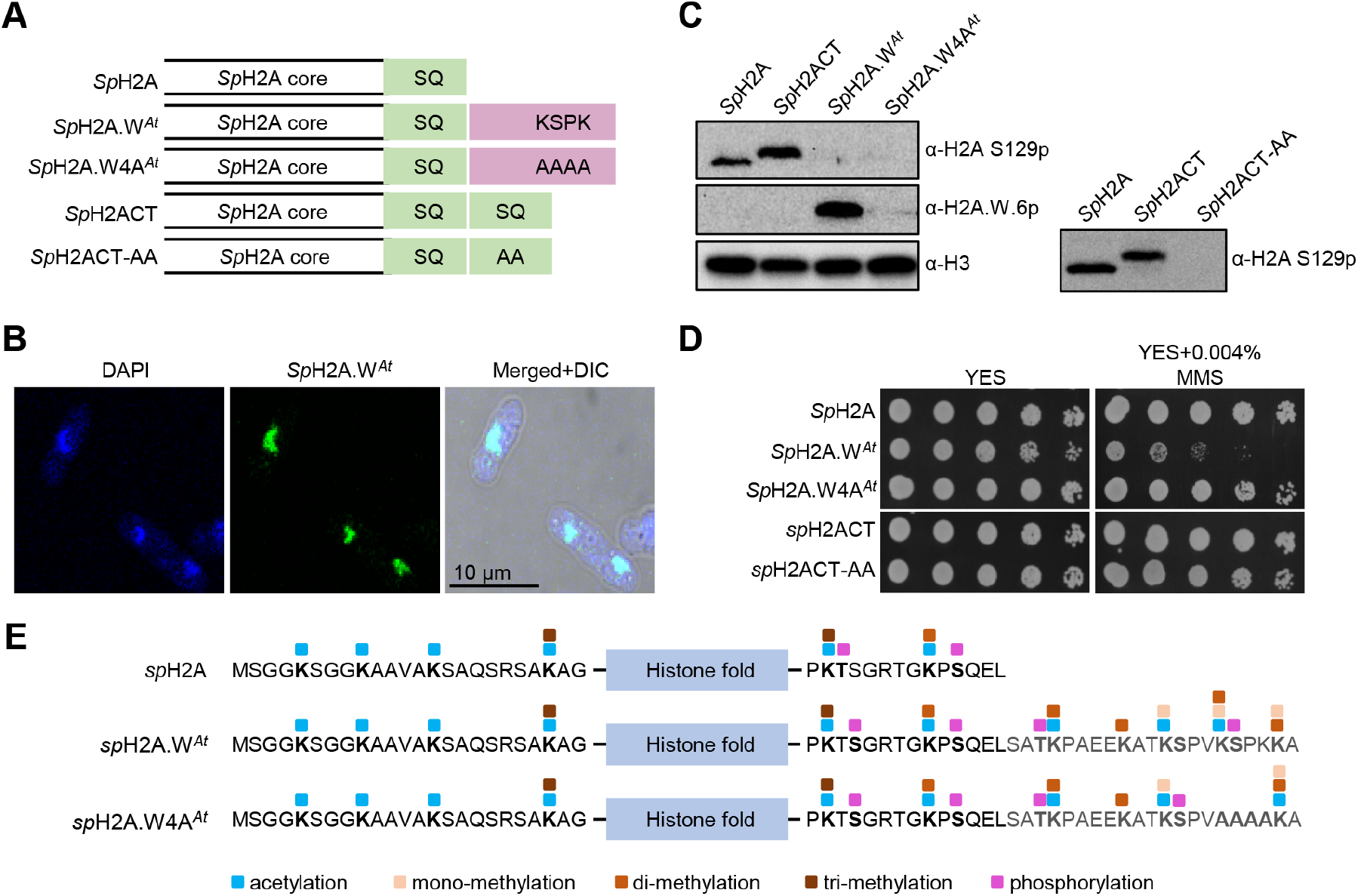
Phosphorylation of the KSPK motifs results in DNA damage sensitivity in yeast. (A) Schematic diagrams of fission yeast histone H2A (*Sp*H2A) and mosaic versions containing either a duplicated C-terminal tail (*Sp*H2ACT) or the C-terminal tail from *Arabidopsis* H2A.W.6 (*Sp*H2A.W^*At*^). Mutant versions in the SQ and KSPK motifs of the latter two constructs, *Sp*H2ACT-AA and *Sp*H2A.W4A^*At*^, are also depicted. (B) *Sp*H2A.W^*At*^ localizes to the nucleus in fission yeast. Confocal images of immunostained cells from exponential phase are shown. 4′,6-diamidino-2-phenylindole (DAPI) staining was used to visualize nuclei and differential interference contrast (DIC) for cell shape. (C) Analysis of *Sp*H2A S129 (SQ) and of *Sp*H2A.W^*At*^ (KSPK) phosphorylation in indicated yeast strains. Cells were collected during exponential growth and whole cell extracts were analyzed with the indicated antibodies, using anti-histone H3 antibody as loading control. The lack of SQ phosphorylation signal in strains expressing *Sp*H2A.W^*At*^, *Sp*H2A.W4A^*At*^, and *Sp*H2ACT-AA is due to the inability of the antibody to bind to the epitope. Phosphorylation of the SQ motif in these strains was confirmed by mass spectrometry (S5D Fig). (D) Phosphorylation of *Sp*H2A.W^*At*^ at the KSPK motif confers sensitivity to DNA damage. Serial dilutions of fission yeast cells expressing WT and indicated *Sp*H2A mosaic variants were spotted onto either YES or YES plates containing 0.004% methyl methane sulfonate (MMS) and incubated at 32°C for 3 days. (E) Summary of all PTMs detected on *Sp*H2A in WT and chimeric histone in indicated yeast strains. Amino acid sequence of N- and C-terminal tails of chimeric histone are indicated with the conserved *Sp*H2A SQ motif and H2A.W.6 KSPK motif respectively. The blue box indicates the histone fold domain. Post-translational modifications detected by MS are color-coded as indicated at the bottom.

The alternative possibility is that phosphorylation of the KSPK motif specifically interfered with DDR events downstream of SQ phosphorylation. We performed an SGA (synthetic gene array) screen with a genome-wide mutant library of non-essential genes to identify candidate genes that display genetic interactions with *Sp*H2A.W^*At*^ in presence of the drug hydroxy urea (HU). From the cluster analysis, cluster 2 contained several groups of genes genetically interacting with *Sp*H2A.W^*At*^, including S/T protein kinases, S/T phosphatases and DNA repair proteins including BRCT domain proteins (Fig S6). We focused on the BRCT domain protein Mdb1 that binds to phosphorylated SQ motif in fission yeast [19]. Previous studies showed that the phosphorylated SQ motif of *Sp*H2A was able to directly bind Mdb1 [19]. To address directly whether presence and/or phosphorylation of KSPK motif interferes with the interaction between Mdb1 and SQ phosphorylation, we performed *in vitro* pull-down assay with synthetic biotinylated peptides (Fig 6A) and recombinant Mdb1. Mdb1 was pulled down by the phosphorylated *Sp*H2A SQ peptide (SpQ) but not by the unmodified version (Fig 6A). Surprisingly, Mdb1 was able to bind the phosphorylated SQ motif in the presence of the KSPK motif but not if this latter motif was phosphorylated (Fig 6A). We concluded that phosphorylation of the KSPK motif specifically interfered with DDR events downstream of SQ phosphorylation and likely prevented binding of Mdb1 to the phosphorylated SQ motif (Fig 6B).

**Fig 6.**
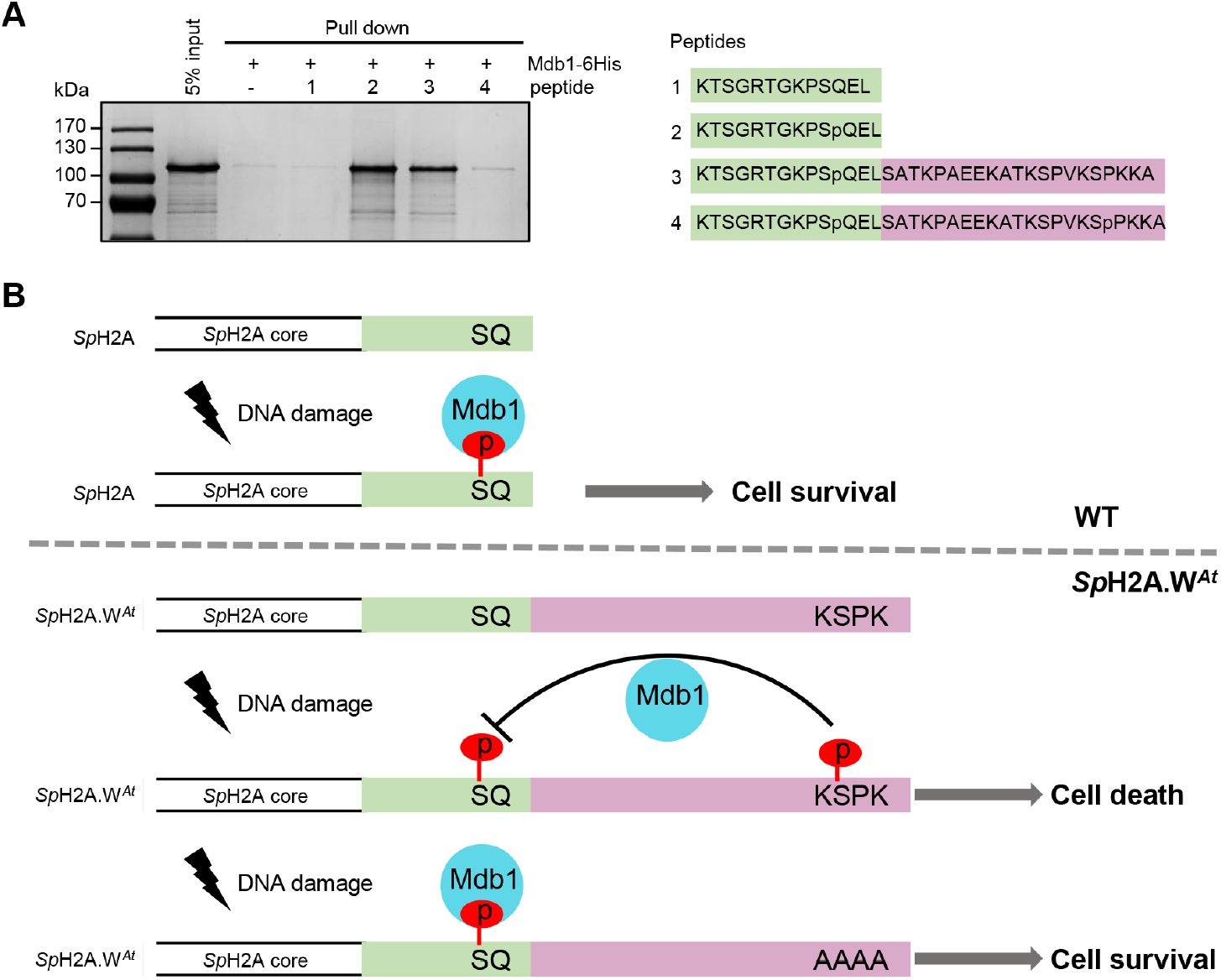
KSPK motif directly interferes with binding of Mdb1 to phosphorylated SQ in a phosphorylation dependent manner. (A) Mdb1 recombinant protein was expressed in *E. coli* BL21 and purified using Ni-NTA beads. Biotinylated peptides, as depicted on the right, corresponding to C terminus of *Sp*H2A, with unmodified (SQ) or S129 phosphorylation (SpQ), or SpQ with unmodified KSPK motif (SpQ+KSPK) or SpQ with S145 phosphorylation (SpQ+KSpPK), were incubated with streptavidin Dynabeads and Mdb1 recombinant protein. The eluates were analyzed by SDS-PAGE with Coomassie staining. (B) The model of crosstalk between SQ and KSPK phosphorylation. In WT cell, S129 phosphorylation site recruit Mdb1 as platform for downstream DDR in response to DNA damage. In *Sp*H2A.W^*At*^ cell, KSPK phosphorylation prevents the Mdb1 binding to SQ phosphorylation site, thus DNA cannot repair properly. In *Sp*H2A.W4A^*At*^ cell, the absence of KSPK phosphorylation allowed the Mdb1 binding to phosphorylated SQ to recruit DDR for DNA repair.

## Discussion

In addition to the well-characterized modifications associated with histone H3, we report a series of modifications at the N- and C-terminal tails, which are either common and specific to the H2A.W family or specific to each isoform. H2A.W.6 is phosphorylated at the serine residue in the KSPK motif in a cell cycle-dependent manner. The kinase CDKA;1, which is active during S phase, is the primary writer of H2A.W.6p, but this mark might also be reinforced by plant specific B-type CDKs active at the G2/M-phase transition [48, 49, 53]. When we compare modifications in H2A variants in *Arabidopsis* with other species, we find that some are plant-specific while others are found in other eukaryotes and show interesting parallels. Like S145 that is specific to H2A.W, S95 is only present in H2A and H2A.X in land plants. Phosphorylation of S95 of the replicative H2A variant has been observed in *Arabidopsis* and implicated in flowering time regulation [56]. Certain isoforms of H2A.W in maize are phosphorylated at S133 during mitosis and meiosis [57]. This residue corresponds to S129 of *Arabidopsis* H2A.W and is absent from H2A.Z, H2A.X, and three out of four H2As. In yeast, the corresponding S121 is phosphorylated by Bub1 to recruit shugoshin, an important step for centromere function in chromosome segregation [58–60]. This serine is replaced by threonine (T120) in human H2A and H2A.X, and its phosphorylation is associated with transcriptional activation and mitotic chromosome segregation [61–63]. Consistent with these data, we detected only a few events of S129 phosphorylation in H2A.W in samples originating from leaves or seedlings that contain low amounts of dividing cells. The neighboring lysine 119 of H2A and corresponding lysines of H2A.X, H2A.Z, and macroH2A are well-known ubiquitination sites, associated with transcriptional repression, gene silencing, and DDR [7, 64–72].

Surprisingly, although KSPK of H2A.W.7 could be phosphorylated *in vitro*, this modification on H2A.W.7 did not occur *in planta*. We showed that prevention of KSPK phosphorylation in H2A.W.7 originates from the sequence of the C-terminal tail of H2A.W.7 and not from its transcriptional pattern. We can speculate that *in planta*, specific modifications of the C-terminal tail of H2A.W.7 prevent the action of the cyclin dependent kinases. We addressed the biological significance of the lack of KSPK phosphorylation on H2A.W.7 *in planta* with a synthetic approach by engineering the endogenous H2A from *S. pombe* to create a histone variant analogous to H2A.W.7. The association of both SQ and KSPK motifs in the C-terminal tail of yeast H2A.X resulted in DNA damage sensitivity, highlighting incompatibility of such association. Results obtained *in vitro* indicate that the phosphorylation of the KSPK (S145) motif prevents BRCT domain protein Mdb1 binding to γH2A.X in response to DNA damage. Transgenic *Arabidopsis* plants expressing combinations of phosphomimetic mutants in the SQ and KSPK motifs further supported this conclusion. We propose that phosphorylation of S145 KSPK prevents the recognition of the phosphorylated SQ motif of H2A by an ortholog of Mdb1, which is still unknown in plants. Recruitment of the SQ motif occurred in members of the H2A.W family in various unrelated species during the evolution of flowering plants, suggesting that the mechanism that we describe in *Arabidopsis* has evolved several times and uses pre-existing components from the DDR and cell cycle.

Altogether our data illustrate that H2A variants extend the histone modification repertoire across eukaryotes beyond that of the extensively studied H3. Importantly, because H2A strongly impact nucleosome and chromatin properties, H2A variant specific modifications and their interplay extend the potential to modulate these properties, adding a new important layer of epigenetic complexity. Furthermore, through the example of interference between PTMs of motifs of H2A.X and of H2A.W, our findings provide a clue to explain why C-terminal motifs that identify classes of H2A variants are mutually exclusive. We propose that PTMs of these motifs mediate all or part of the function of the motifs themselves. The presence of two motifs side by side might interfere with the function of each motif, leading the emergence of distinct classes of H2A variants differentiated by specific functional motifs with distinct biological functions.

## Methods

### Plant material

The mutant lines *hta6* (SALK_024544.32; *h2a*.*w*.*6*), *hta7* (GK_149G05; *h2a*.*w*.*7*), and *hta12* (SAIL_667_G10; *h2a*.*w*.*12*) have been published [21]. Transgenic lines expressing WT and hypomorphic SQ motif mutants of H2A.W.7 in the *hta7* mutant background have been published [16]. The hypomorphic CDKA;1 D and DE mutants [51, 52] and transgenic line expressing CDKA;1-YFP in *cdka;1* mutant background [73] were a kind gift from Dr. Arp Schnittger. Primers used for genotyping hypomorphic CDKA;1 D and DE mutants (S4C Fig) were: Tg9WT/TDNA_N049 (TGTACAAGCGAATAAAGACATTTGA), Tg9WT_N048 (CAGATCTCTTCCTGGTTATTCACA), and Tg9TDNASalk_LB_J504 (GCGTGGACCGCTTGCTGCAACTCTCTCAGG). Unless otherwise stated, plants were grown in a fully automated climate chamber at 21°C, under long day conditions.

To construct mutant versions of H2A.W.7 containing either WT SQ motif or mutated SQ motif into AQ in combination with phosphomimic and nonphosphorylatable versions of the KSPK motif (KDPK or KAPK, respectively), a two-step PCR reaction was performed using previously described constructs expressing H2A.W.7 containing either the WT SQ or mutated AQ motif [16] as templates. For the first step amplification, the following primer pairs were used: W7-fw (GTACTCTAGAGAGGCATAGATACCGCGCCATC) with either W7-Rev1KDP (AGGATCTTTGGTAGCAGAAG) or W7-Rev1KAP (AGGAGCTTTGGTAGCAGAAG). The second PCR step used the W7-Rev2KDP (AGCAGGATCCAGCCTTCTTAGGATCTTTGGT) or W7-Rev2KAP (AGCAGGATCCAGCCTTCTTAGGAGCTTTGGT) as reverse primers with the same forward primer as above. The mutant *HTA7* fragments were cut with *Bam*HI and *Xba*I and ligated into pCBK02. *hta7* mutant plants were transformed by the floral dip method and transgenic plants were selected on MS plates containing 10 µg/ml of phosphinothricin. The T3 and T4 homozygous seeds containing single transgene copy were used for experiments. For swapping the promotors and tails of H2A.W.6 and H2A.W.7, cloning primers (S1 Table) were used to amplify the different fragments from genomic DNA of *Arabidopsis* seedlings. We created the green gate entry modules following previously published protocol [74]. The promotor fragments were introduced into the A-B empty vector, the full H2A.W.6 and H2A.W.7 as well as the N-terminal+core fragments were introduced into the C-D empty vector and the tail fragments were introduced into the D-E empty vectors. The Ub10 Terminator in the E-F entry vector and the selection based on seed coat YFP fluorescence were a gift from Yasin Dagdas laboratory. For the final greengate reaction pGGZ003_ccdB, the entry clones were combined with empty destination vector. The final constructs were transformed into the triple *hta6 hta7 hta12* mutant which is a complete knock-out line deprived of H2A.W described in the preprint doi: https://doi.org/10.1101/2020.03.19.998609.

### Isolation of nuclei, micrococcal nuclease (MNase) chromatin digestions, immunoprecipitation, and SDS-PAGE

Nuclei isolation, MNase digestion, and immunoprecipitation were performed as previously described [16] by using 4 grams of 2-3 week old leaves for each antibody. Isolated nuclei were washed once in 1 ml of N buffer (15 mM Tris-HCl pH 7.5, 60 mM KCl, 15 mM NaCl, 5 mM MgCl_2_, 1 mM CaCl_2_, 250 mM sucrose, 1 mM DTT, 10 mM ß-glycerophosphate) supplemented with protease and phosphatase inhibitors (Roche). After centrifuging for 5 min at 1,800 × *g* at 4°C, nuclei were re-suspended in N buffer to a volume of 1 ml. Twenty µl of MNase (0.1 u/µl) (SigmaAldrich) were added to each tube and incubated for 15 min at 37°C. During the incubation, nuclei were mixed 4 times by inverting the tubes. MNase digestion was stopped on ice by the addition of 110 µl of MNase stop solution (100 mM EDTA, 100 mM EGTA). Nuclei were lysed by the addition of 110 µl of 5 M NaCl (final concentration of 500 mM NaCl). The suspension was mixed by inverting the tubes and they were then kept on ice for 15 min. Extracts were cleared by centrifugation for 10 min at 20,000 × *g* at 4°C. Supernatants were collected and centrifuged again as above. For each immunoprecipitation extract, an equivalent of 4 g of leaf material was used, usually in a volume of 1 ml. To control MNase digestion efficiency, 200 µl of the extract were kept for DNA extraction. Antibodies, including non-specific IgG from rabbit, were bound to protein A magnetic beads (GE Healthcare) and then incubated with MNase extracts over night at 4°C. Beads were washed 2 times with N buffer without sucrose containing 300 mM NaCl, followed by 3 washes with N buffer containing 500 mM NaCl without sucrose, and 1 wash with N buffer without sucrose, containing 150 mM NaCl. Beads were incubated 2 times with 15 µl of hot loading buffer for 5 min and supernatants were removed and combined. Proteins were resolved on 4-20% gradient gels (Serva) and silver stained.

### Generation of antibodies, isolation of nuclei, SDS-PAGE, and western blotting

Antibodies against H2A.X, H2A.W.6, H2A.W.7, γH2A.X, and γH2A.W.7 have been described [16, 21]. Antibodies against the H2A.W.6 phosphopeptide (CEEKATKSPVKSpPKKA) were raised in rabbits (Eurogentec) and purified by peptide affinity column. Purified IgG fractions were tested for specificity on peptide arrays containing serial dilutions of non-phosphorylated and phosphorylated peptides (phospho-serine specific antibodies).

Nuclei for western blot analyses were prepared from 300 mg of tissue (10-12 day old treated seedlings) as described [16]. Tissue was frozen in liquid nitrogen and disrupted in 15 ml Falcon tubes by rigorous vortexing with 5 small ceramic beads. Ten ml of nuclei isolation buffer (NIB; 10 mM MES-KOH pH 5.3, 10 mM NaCl, 10 mM KCl, 250 mM sucrose, 2.5 mM EDTA, 2.5 mM ß-mercaptoethanol, 0.1 mM spermine, 0.1 mM spermidine, 0.3% Triton X-100), supplemented with protease and phosphatase inhibitors (Roche) were added followed by short vortexing to obtain a fine suspension. The suspension was filtered through 2 layers of Miracloth (Merck Millipore) into 50 ml Falcon tubes, followed by washing the Miracloth with NIB. The sample was centrifuged at 3,000 rpm for 5 min at 4°C. The pellet was resuspended in NIB buffer and transferred into an Eppendorf tube. The sample was centrifuged for 2 min at 4°C at full speed and the resulting pellet was resuspended in 0.3 × PBS in 1 × Laemmli loading buffer. The sample was then boiled for 5 min and immediately centrifuged for 2 min at maximum speed to remove starch and other large particles. For western blot analyses, 10 µl for histone variants, 20 µl for histone modifications, and 10 µl for H3, used as a loading control, were loaded per lane. Western blots with phospho-specific antibodies were performed in solutions containing TBS instead of PBS and, after blocking, the membranes were incubated with primary antibody in TBS without milk.

Antibodies against H2A.X, H2A.W.6, H2A.W.7, γH2A.X, and γH2A.W.7 [16, 21], as well as H2A.W.6p (this work), H3 (ab1791, Abcam), CDK (PSTAIR, Sigma P7962), and CDKY15p (Cell Signaling Technology) were used at 1:1,000 dilution and secondary antibodies, HRP conjugated goat anti-rabbit and goat anti-mouse, at 1:10,000. Signals were detected using the chemiluminescence kit (ThermoFisher), recorded using an ImageDoc instrument (BioRad), exported to Photoshop, and prepared for publication. Quantification of western blots was done with ImageLab 5.2 software (BioRad) using the volume tool. For detection of Y15 phosphorylation of CDK, the manufacturer’s protocol was followed.

### Nano LC-MS/MS analysis

Histone bands corresponding to H2A.W.6/H2A.W.7 from *Arabidopsis* and *Sp*H2A/*Sp*H2A.W/*Sp*H2A.W4A from fission yeast were excised from silver stained gels, reduced, alkylated, in-gel trypsin, LysC, and subtilisin digested, and processed for MS. The nano HPLC system used was a Dionex UltiMate 3000 HPLC RSLC (Thermo Fisher Scientific, Amsterdam, Netherlands) coupled to a Q Exactive HF mass spectrometer (Thermo Fisher Scientific, Bremen, Germany), equipped with a Proxeon nanospray source (Thermo Fisher Scientific, Odense, Denmark). Peptides were loaded onto a trap column (Thermo Fisher Scientific, Amsterdam, Netherlands, PepMap C18, 5 mm × 300 μm ID, 5 μm particles, 100 Å pore size) at a flow rate of 25 μl/min using 0.1% TFA as the mobile phase. After 10 min, the trap column was switched in line with the analytical column (Thermo Fisher Scientific, Amsterdam, Netherlands, PepMap C18, 500 mm × 75 μm ID, 2 μm, 100 Å). Peptides were eluted using a flow rate of 230 nl/min, and a binary 1-hour gradient.

The Q Exactive HF mass spectrometer was operated in data-dependent mode, using a full scan (m/z range 380-1500, nominal resolution of 60,000, target value 1E6) followed by MS/MS scans of the 10 most abundant ions. MS/MS spectra were acquired using normalized collision energy of 27%, isolation width of 1.4 m/z, resolution of 30,000 and the target value was set to 1E5. Precursor ions selected for fragmentation (exclude charge state 1, 7, 8, >8) were put on a dynamic exclusion list for 20 s. Additionally, the minimum AGC target was set to 5E3 and intensity threshold was calculated to be 4.8E4. The peptide match feature was set to preferred and the exclude isotopes feature was enabled. For peptide identification, the RAW files were loaded into Proteome Discoverer (version 2.1.0.81, Thermo Scientific). All hereby created MS/MS spectra were searched using Mascot 2.2.7 against a database which contains all histone variants from *Arabidopsis thaliana*. The following search parameters were used: Beta-methylthiolation on cysteine was set as a fixed modification, oxidation on methionine, deamidation on asparagine and glutamine, acetylation on lysine, phosphorylation on serine, threonine and tyrosine, methylation and di-methylation on lysine and arginine and tri-methylation on lysine were set as variable modifications. Monoisotopic masses were searched within unrestricted protein masses for tryptic enzymatic specificity. The peptide mass tolerance was set to ±5 ppm and the fragment mass tolerance to ±0.03 Da. The result was filtered to 1% FDR at the peptide level using the Percolator algorithm integrated in Thermo Proteome Discoverer and additional a minimum Mascot score of 20. In addition, we have checked the quality of the MS/MS spectra manually. The localization of the phosphorylation sites within the peptides was performed with ptmRS using a probability threshold of minimum 75 [75].

### DNA damage sensitivity assays

For the true leaf assay, sterilized seeds were put on MS plates containing 50 µg/ml of zeocin (Invitrogen) and, after stratification at 4°C for 3 days, germinated under long day conditions. Development of true leaves was scored 12 days after germination. For analysis of H2A.W.7, H2A.X, and H2A.W.6 phosphorylation in response to DNA damage, 300 mg of twelve-day old seedlings germinated and grown on MS plates under long day conditions were treated in liquid MS medium with 20 µg/ml bleomycin (Calbiochem) for the indicated time periods or for 2 hours with 20 µg/ml bleomycin. After treatment, seedlings were removed from medium and shock frozen in liquid nitrogen for nuclei isolation and western blot analysis as described above.

### Cell cycle synchronization of BY-2 cell suspension culture with aphidicolin

The BY-2 cell suspension culture was sub-cultured every 2 weeks by adding 1 ml of the previous culture to 50 ml fresh BY-2 media. The culture was kept in the dark and under constant shaking at 130 rpm at room temperature. For cell cycle synchronization, a previously published protocol was followed [76]. In short, 1.5 ml of stationary BY-2 cells were added to 95 ml of fresh BY-2 media and grown for 5 days at 27°C. This culture was diluted 4 times with fresh BY-2 media and then treated with 20 µg/ml aphidicolin (1 mg/ml; SigmaAldrich) for 24 hours at 27°C to reach a cell cycle block. To release the cells from this block, the cells were passed over a 40 µm sieve, washed with fresh media, and resuspended in 100 ml of fresh media and followed in a time course of 12 hours. Every hour, a sample was taken for western blotting and flow cytometry analysis. For western blotting, 5 ml of cells were collected with a 50 µm sieve and immediately frozen in liquid nitrogen. To extract proteins, samples were crushed with a small pistil in an Eppendorf tube and mixed with 200-250 µl of 1 × loading buffer in 1 × PBS. Samples were boiled for 5 min at 99°C followed by centrifugation at full speed for 5 min. Fifteen µl were loaded onto 15% SDS-PAGE gels and analyzed by western blotting.

For the flow cytometry analysis, 2 ml of cells were collected with a 50 µm sieve. The cells were resuspended in a small Petri dish in 400 µl of extraction buffer (CyStain UV Precise P kit from Sysmex) and chopped with a razor blade. The sample was transferred onto a 50 µm CellTrics Disposable filter (Partec), placed on top of a flow cytometry tube, and 1,5 ml of DAPI stain solution (CyStain UV Precise P kit from Sysmex) was added to the sieve. The samples were then analyzed by the Partec flow cytometer with a gain set to 380.

### Staining of *Arabidopsis* roots with EdU

For the whole mount EdU staining and immunostaining with the phosphorylation specific antibody in roots, the protocol for the Click-iT EdU imaging kit from Invitrogen was combined with the protocol from [77]. *Arabidopsis* plants were grown on MS plates for 1 week and fifteen seedlings were incubated for 1 hour in liquid MS media containing 10 µM EdU at room temperature. The seedlings were washed twice with MS media to remove the EdU. The roots were cut off and transferred into an Eppendorf tube with fixative solution, incubated for 1 hour at room temperature, and washed twice for 10 min with 1×PBS and twice for 5 min with water. The roots were transferred to the microscope slide and allowed to dry. The material was then rehydrated by adding 1 × PBS for 5 min followed by incubation with 2% driselase (Sigma) for 60 min at 37°C in a moisture chamber to remove the cell wall. The roots were washed 6 times for 5 min with 1×PBS at room temperature. Mixture of 3% IGEPAL CA-630 plus 10% DMSO was added to the slide and incubated for 60 min at room temperature. The solution was removed, and the slides were washed 8 times with 1 × PBS for 5 min. The slides were then incubated with 3% BSA in 1 × PBS for 5 min before revealing EdU incorporation with the Click-iT reaction buffer. After this reaction, the slides were washed once for 5 min with 3% BSA in 1 × PBS followed by blocking for 60 min at room temperature with 3% BSA in 1 × PBS. Approximately 150 µl of the primary antibody diluted 1:100 in 3% BSA 1 × PBS was added to the slides and incubated for 4 hours at 37°C in a humid chamber and then overnight at 4°C. Before incubating with the secondary antibody, the slides were washed 5 times with 1 × PBS for 10 min at room temperature. Alexa flour 555 labelled goat anti-rabbit IgG (Life Technologies) diluted 1:200 in 1 × PBS containing 3% BSA was added to slides and incubated for 3 hours at 37°C in a humid chamber. Finally, slides were again washed 5 times for 10 min with 1 × PBS at room temperature and mounted in Vectashield (Vector Laboratories) with 1 µg/ml DAPI. Slides were examined on an LSC confocal microscope (Carl Zeiss) and confocal sections were acquired with a 40 × oil objective, exported to Adobe Photoshop, and prepared for publication.

### Treatment of *Arabidopsis* cell suspension with cell cycle inhibitors

The T87 cell suspension was grown on a shaker at 130 rpm under constant light. For treatment with different cell cycle inhibitors, 5 ml of the 7-day old culture were mixed with 5 ml of fresh BY-2 media supplemented with the indicated amounts of the inhibitors. The concentrations that were used for the treatment were: 50 µM roscovitine (50 mM solution from Merck); 10 mM hydroxyurea (SigmaAldrich); 4 µg/ml aphidicolin (SigmaAldrich). Cells were incubated with the inhibitors for 24 hours and subsequently collected by removing the media with a 50 µm mesh, shock frozen in liquid nitrogen, and stored at −80°C.

For treatment of cell suspension cultures with different concentrations of roscovitine, a 7-day old *Arabidopsis* T87 culture was diluted in BY-2 media to an OD_600_ of 0.167. 10 ml of the diluted culture were incubated for 24 hours with or without 10 µM, 20 µM, 30 µM, 40 µM, and 50 µM roscovitine with shaking under constant light. Seven ml of each sample were collected, and the cells were disrupted with ceramic beads and immediately mixed with 1 × Laemmli loading buffer in 0.3 × PBS without prior enrichment of nuclei.

### Immunoprecipitation of CDKA;1 from *cdka-/-* CDKA;1::YFP plants

For immunoprecipitation, 1.5 g of 15 day old CDKA;1::YFP *cdka-/-* [73] and WT seedlings grown on MS plates were crushed in liquid nitrogen and powder was resuspended in PEB400 buffer (50 mM HEPES-KOH pH 7.9, 400 mM KCl, 1 mM DTT, 2.5 mM MgCl_2_, 1 mM EDTA, 0.1% Triton X-100) [78] supplemented with protease inhibitor cocktail (Roche) (100 µl of buffer per 100 mg of seedlings) and the suspension was incubated on ice for 10 minutes. The samples were centrifuged for 10 min at full speed at 4°C. The supernatants were transferred to a new tube and centrifuged as above. The volume of the sample was measured and the same volume of PEB buffer without KCl was added to obtain the PEB buffer with 200 mM KCl. At this step, an input aliquot was taken and mixed with 5 × loading buffer. Agarose beads coupled with GFP nanobodies (40 µl; VBC Molecular Biology Services) were washed twice with PEB200 and the protein extracts were added to the beads and the mixture was incubated overnight at 4°C on a rotating wheel. The beads were washed 3 times with 1 ml of PEB200 followed by 2 washes with kinase buffer (20 mM Tris-HCl pH7.5, 50 mM KCl, 5 mM MgCl_2_, 1 mM DTT) if they were used for the *in vitro* kinase assay. To analyze immunoprecipitations by SDS-PAGE and western blotting, 30 µl of 1 × loading buffer in PEB200 were added to the beads. After boiling for 5 min, the supernatants were loaded onto a 12% SDS-PAGE gel.

### *In vitro* phosphorylation assay

For the kinase assay, 10 µl of kinase buffer were added to the agarose beads with the immunoprecipitated CDKA;1-YFP. One µg of the reconstituted histone dimers (H2A.W.6-H2B.9, H2A.W.7-H2B.9, H2A.13-H2B.9) and 200 µM ATP were added and the reaction was mixed before incubating for 35 min at 30°C. As a control, 20 µl kinase buffer were mixed with the dimers and ATP for 35 min at 30°C. Reactions were stopped by adding 5× loading buffer and analyzed by SDS-PAGE followed by western blotting with H2A.W.6p antibody.

### Cloning of H2A.W.7 into overexpression vector pET15b

cDNA encoding H2A.W.7 was PCR amplified from a seedling cDNA library using the following primers: W7pET15bfor (GCATCCATATGGAGTCATCACAA) and W7pET15brev (CTAATGGATCCTCAAGCCTTCTT). The PCR product was cleaned using the PCR purification kit from Qiagen, digested with *Nde*I/*Bam*HI, gel purified, and ligated into *Nde*I/*Bam*HI opened pET15b (Novagene). Plasmids for expression of His-tagged H2A.W.6, H2A.13, and H2B.9 have been published [21, 22].

### Overexpression and purification of recombinant histones and assembly of H2A-H2B dimers

Proteins were overexpressed in *E. coli* BL21 (DE3) overnight at 37°C. Histone purification was performed as previously described [79, 80]. Cells pellets were resuspended in sonication buffer 1 (50 mM Tris-HCl pH8.0, 500 mM NaCl, 1 mM PMSF, 5% glycerol) and sonicated with 50% power for 5 min. After centrifugation at 15,000 rpm at 4°C for 20 min, pellets were resuspended in sonication buffer 1 and sonicated at 35% power for 2 min followed by centrifugation as before. The resulting pellets were resuspended in sonication buffer 2 (50 mM Tris-HCl pH 8.0, 500 mM NaCl, 7 M guanidine hydrochloride, 5% glycerol) and sonicated as described before. After the third sonication, the suspension was rotated at 4°C overnight. After centrifugation, the supernatants containing denatured proteins were mixed with Ni-NTA resin (Qiagen) and incubated for 60 min at 4°C. To remove the supernatant, the suspension was centrifuged. The resin was resuspended in wash buffer (50 mM Tris-HCl pH8.0, 500 mM NaCl, 6 M urea, 5% glycerol, 5 mM imidazole) and transferred into an Econo-column (BioRad). After washing with 50 column volumes of wash buffer, proteins were eluted with elution buffer (50 mM Tris-HCl pH8.0, 500 mM NaCl, 6 M urea, 5% glycerol, 500 mM imidazole) and collected in 2 ml fractions. The fractions were analyzed on a 15% SDS-PAGE gel and those containing histones were pooled together and dialyzed against 4L of 10 mM Tris-HCl pH 7.5, 2 mM 2-mercaptoethanol at 4°C for 2 days. After checking the thrombin cleavage efficiency with different U/mg of thrombin, the estimated amount of thrombin was added to each sample and incubated for 3 hours at room temperature followed by analysis on a 15% SDS-PAGE gel. Proteins were further purified by cation ion exchange chromatography. The sample was applied to an SP sepharose column (GE Healthcare) connected to an Äkta system. The column was equilibrated and washed with equilibration buffer (20 mM CH_3_COONa pH5.2, 200 mM NaCl, 6 M urea, 5 mM 2-mercaptoethanol, 1 mM EDTA). For elution, a linear gradient of 200-800 mM NaCl in elution buffer (20 mM CH_3_COONa pH 5.2, 6 M urea, 5 mM ß-mercaptoethanol, 1 mM EDTA) was used. The fractions were again analyzed on a 15% SDS-PAGE gel. Histone containing fractions were pooled together and dialyzed against 4 L of 2 mM 2-mercaptoethanol 4 times for 4 hours. Finally, the purified histones were freeze-dried.

For histone H2A-H2B dimer reconstitution, freeze-dried histones were resolved in unfolding buffer (20 mM Tris-HCl pH 7.5, 7 M guanidine hydrochloride, 20 mM 2-mercaptoethanol) at a concentration of 1 mg/ml at a 1:1 molar ratio and incubated for 2 hours at 4°C on a wheel. Samples were step-wise dialyzed against refolding buffer (20 mM, Tris-HCl pH 7.5, 1 mM EDTA, 0.1 M PMSF, 5%glycerol, 5 mM 2-mercaptoethanol) staring with 2 M NaCl at 4°C overnight, followed by 1 M NaCl refolding buffer at 4°C for 4 hours, 0.5 M NaCl refolding buffer at 4°C for 4 hours, and finally against 0.1 M NaCl refolding buffer at 4°C overnight. Proteins were concentrated by centrifugation with a 10 kDa cut-off membrane (Merck Millipore) to a volume of 300 µl. The sample was applied to a Superdex200 gel filtration column in 0.1 M NaCl refolding buffer. Peak fractions were analyzed on a 15% SDS-PAGE gel and fractions containing the heterodimers were pooled together. The heterodimers were then concentrated by ultrafiltration (10 kDa cut-off) and, at the same time, the buffer was exchanged to kinase buffer. The final protein concentration was determined by measuring the absorbance at 280 nm and the quality was analyzed on a 15% SDS-PAGE gel.

### *Schizosaccharomyces pombe* strains and media

Unless otherwise stated, cells were grown in rich medium (YES). Gene replacement and tagging were performed by homologous recombination using a plasmid-based method [81]. The primers used are listed in S1 Table. In brief, pCloneNat1 and pCloneHyg1 were used to fuse the H2A.W.6 (^*At*^) C-terminal 21 amino acids to the endogenous C-terminus of *Sp*H2A.α and *Sp*H2A.β by using the natMX4 or hphMX4 cassette, respectively [25]. All constructed strains in this study were verified by PCR analysis and sequencing.

### *S. pombe* chromatin fractionation assay

Chromatin fractionation of WT and *Sp*H2A.W^*At*^ strains was performed as previously described [82] and fractions were analyzed by western blotting using anti-αtubulin (T6199, Sigma), anti-H3 (ab1791, Abcam), and anti-H2A.W.6 (antibodies recognizing C-terminal KSPKKA motif).

### Preparation of *S. pombe* whole cell extracts and acid extraction of histones for MS analysis

Cells were disrupted by 0.5 mm glass beads in lysis buffer (50 mM Tris-HCl pH 7.5, 150 mM NaCl, 5 mM EDTA, 10% glycerol and 1 mM PMSF) and centrifuged at 14,000 × *g* for 15 min at 4°C. Supernatant was collected as a whole cell extract. Histones were analyzed by western blotting with antibodies against fission yeast-specific phosphoS129 of H2A (ab17353, Abcam), H2A.W.6p, and H3 (ab1791, Abcam). For mass spectrometry, histones from WT, *Sp*H2ACT, *Sp*H2ACT-AA, *Sp*H2A.W^*At*^, and *Sp*H2A.W4A^*At*^ cells were prepared by the acid extraction and acetone precipitation method [83].

### Indirect immunofluorescence on *S. pombe* cells

For detection of *Sp*H2A.W^*At*^ by immunofluorescence, cells expressing FLAG-tagged *Sp*H2A.W^*At*^ were fixed with 4% paraformaldehyde, digested to spheroplasts with zymolyase (Zymo Research), permeabilized with 1% Triton X-100, and incubated with an anti-Flag antibody (F1804, Sigma) as the primary antibody at a 1:100 dilution and anti-mouse IgG Alexa Fluor 488 as the secondary antibody (A11029, Life Technologies) at a 1:100 dilution. Cells were placed onto poly L-lysine-coated coverslips and DAPI stained. Microscopic analysis was performed using LSM700 laser scanning confocal microscope (Zeiss).

### Sensitivity of *S. pombe* strains to MMS

Cells were grown in YES medium at 32°C until the exponential phase and for all strains the OD_600_ was adjusted to 1.0. Five-fold serial dilutions were made with fresh YES and 2 μl were spotted on YES plates or YES plates containing 0.004% methyl methane sulfonate (MMS) and incubated at 32°C for 3 days.

### Peptide Pull Down assay

6His-tagged Mdb1 recombinant protein was expressed in BL21 and purified using Ni-NTA beads (Qiagen) following manufacturer’s instructions. As control, we synthesized two peptides corresponding to residues 120–132 of yeast H2A C-terminal tail, with serine 129 being unphosphorylated (termed as SQ) and phosphorylated (termed as SpQ). The other two peptides corresponding to residues 129–149 of H2A.W6 from *Arabidopsis*, with serine 145 being unphosphorylated (termed as KSPK) and phosphorylated (termed as KSpPK) together with SpQ peptide. All these peptides coupled with biotin at N-terminal. Twenty micrograms of each peptide were incubated with 20 µl of pre-washed Dynabeads M-280 Streptavidin (Invitrogen) at RT for 1 hour, then incubated with purified Mdb1 in peptide binding buffer (50 mM Tris-HCl, pH 7.5, 100 mM NaCl, 0.05% NP-40) at 4°C for overnight. Beads were washed with peptide binding buffer and eluted with SDS loading buffer.

### Yeast Genetic interaction screening

Large-scale crosses by SGA (synthetic genetic array) were carried out using the Bioneer haploid deletion mutant library (version 3.0) and the query strains WT and *sp*H2A.W^*At*^. Manipulations were performed using a Singer RoToR colony pinning robot, essentially as described previously [84]. First, the library and query strains were arrayed in 384 colony format on YES agar containing 250 µg/ml G418 (Geneticin; Life Technologies, 10131027) or 100 µg/ml ClonNat (Nourseothricin; Jena Bioscience, AB-102XL) and 50 µg/ml Hygromycin B (Invitrogen/Life Technologies, 10687010), respectively. Mating between library and query strains, and selection of progeny was largely performed as previously reported [85]. Briefly, query and library stain cells were mixed in a drop of sterile H_2_O on SPAS plates and allowed to sporulate at 24°C for 3 days. The resulting cell/spore mixture was incubated at 42°C for 7 days to enrich for spores and plated onto selective media to select for haploid progeny bearing the gene deletion and/or *sp*H2A.W^*At*^. Finally, the selected haploid progeny was grown YES or YES supplemented with or 9 mM hydroxyurea (Sigma, H8627) at 32°C. For each individual screen with the WT and *sp*H2A.W^*At*^ query strains, technical duplicates were processed following the germination step, and each screen was repeated independently at least 3 times, resulting in a total number of n = 6 screen for data analysis.

### Data analysis of the genetic screen and visualization

SGA analysis was performed as previously described [86], with some modifications. Genetic interactions were assessed by colony growth on YES plates containing no additives (non-selective media; N/S) and YES plates containing formamide or hydroxyurea (treatment) and quantified by determining colony sizes (area) of digitalized pictures. As colonies at the edges of the plates show increased growth due to better availability of nutrients, the size of colonies of the first and second outer most rims was corrected by multiplying with a correction factor. For calculating this factor, we determined the ratio between the median of all colonies in the middle of the plate (i.e., excluding the first and second rims) and the median of the first and second outer most rims. Next, the sensitivity towards stress conditions was determined by calculating the ratio between growth on ‘treatment’ and ‘N/S’ for each individual mutant; the obtained value for each mutant was then normalized to the median of the treatment: N/S values derived from all mutants of the same 384-plate. Finally, the entire dataset of each screen was then log_2_-transformed and median-normalized to all mutants from the entire screen (array). To select for robust genetic interactions, data from all screens, query strains and stress conditions were filtered to select for those mutants containing at least 80% of present values and show at least 5 absolute values higher or equal to 0.3 (log_2_), resulting in the selection of 529 mutants. Next, hierarchical clustering of the genetic interaction dataset was performed using Euclidean distance as the similarity metric and complete linkage as the clustering method using Gene Cluster 3.0. Hierarchical cluster maps were visualized with Java TreeView (version 1.1.6) with negative and positive values represented in blue and yellow, respectively (grey: missing or N/A data).

## SUPPLEMENTARY DATA

Supplementary Data are available at Journal Online.

## Authors contributions

AS, BL and ZL performed experiments, and contributed data. FB, ZL, AS and BL interpreted data and wrote the manuscript. KM contributed Mass spectrometry analysis. MC and SB contributed the genetic screen and participated in revising the manuscript. PB and OM contributed the triple mutant line deprived of H2A.W. FB KM and SB were responsible for supervision.

## Acknowledgments

We thank technical assistance from the Vienna Biocenter Core Facilities for Plant Science and the IMP/IMBA/GMI BioOptics and Protein Chemistry facilities for support (Ines Steinmacher, Susanne Opravil, Gabriela Krssakova, Otto Hudecz and Richard Imre).

## Supplemental Figures

**S1 Fig.**
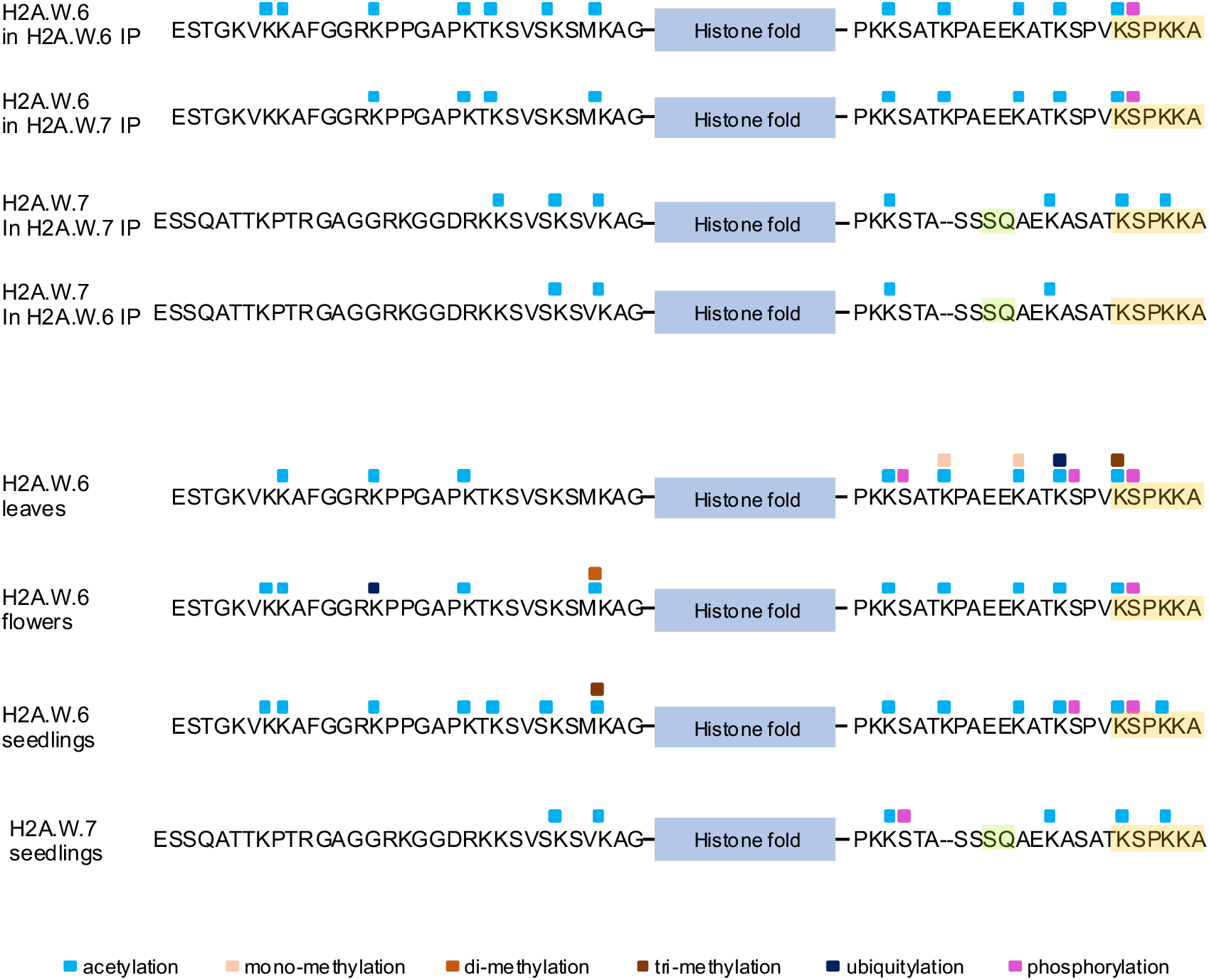
Mass spectrometry analysis of H2A.W.6 and H2A.W.7 modifications. Summary of all modifications detected with samples from leaves, flowers, and seedlings.

**S2 Fig.**
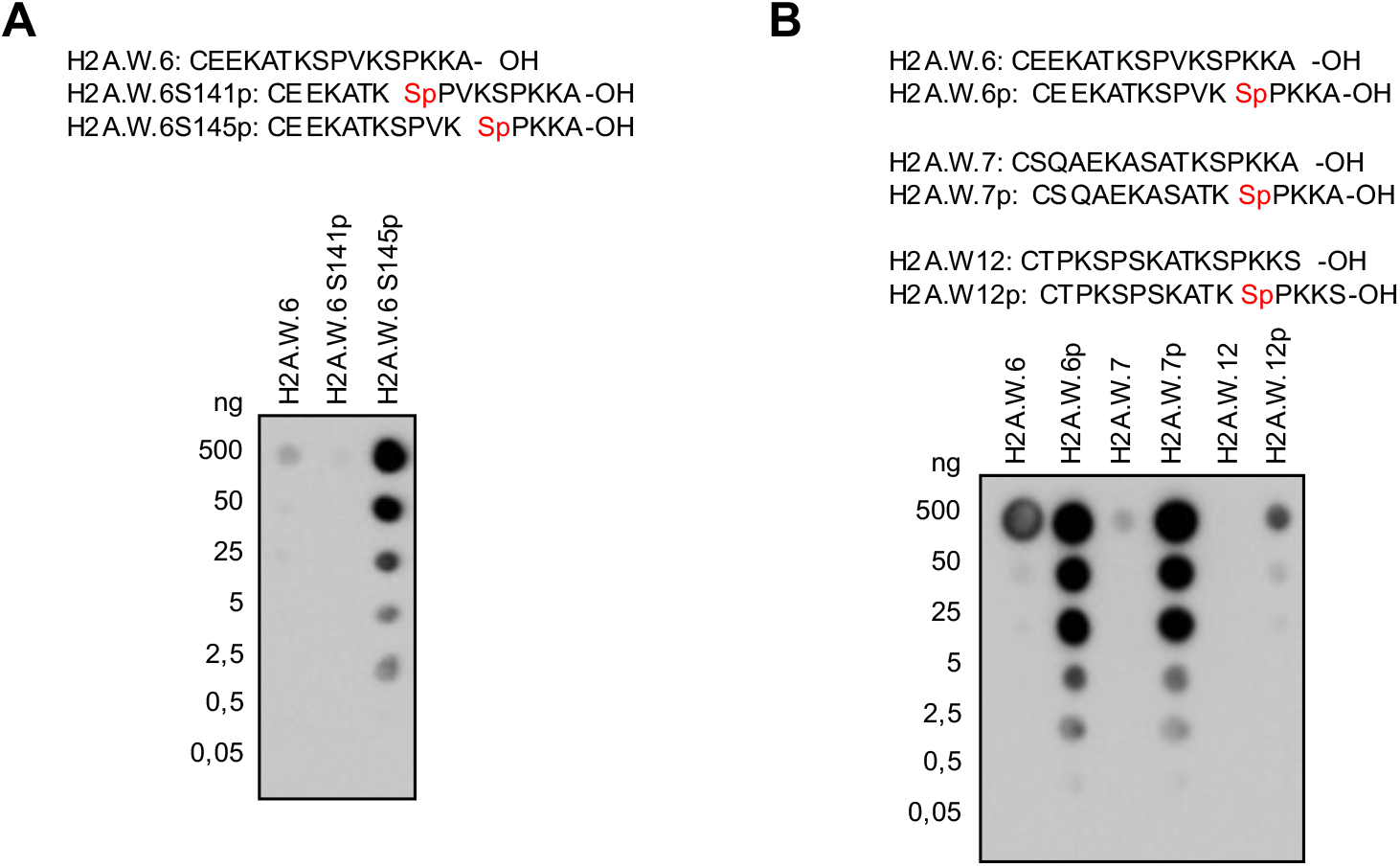
Antibody against the phosphorylated serine 145 of H2A.W.6 is specific but recognizes the phosphorylated KSPKKA motif of H2A.W.7. (**A**) Affinity purified antibody obtained after immunization of rabbits with the CEEKATKSPVKSpPKKA-OH peptide were tested on dot blots with the serially diluted peptides indicated at the top. Note that unphosphorylated peptide or peptide phosphorylated at serine 141 do not cross-react with the affinity purified antibody. (**B**) The same purified antibody was tested against C-terminal peptides in the unphosphorylated and phosphorylated forms from H2A.W.6, H2A.W.7, and H2A.W.12 as above. Note that the H2A.W.7 but not the H2A.W.12 peptide cross reacts with the H2A.W.6p antibody, indicating that the epitope recognized includes the C-terminal alanine that is absent in H2A.W.12.

**S3 Fig.**
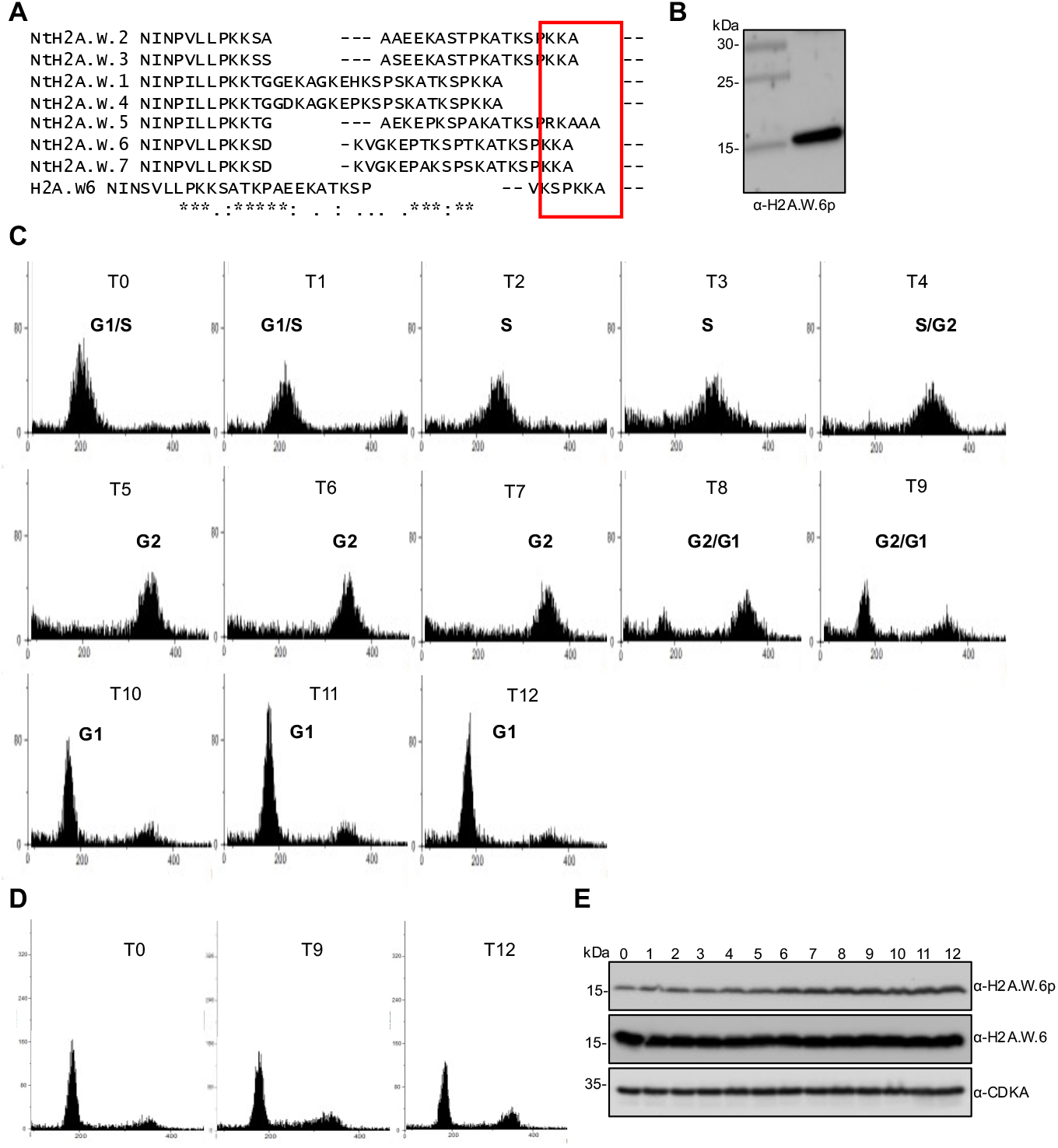
Phosphorylation of the KSPKK motif changes during the cell cycle. (**A**) Protein sequence alignment of tobacco H2A.W isoforms with *Arabidopsis* H2A.W.6. Only the C-terminal tail is shown, with the conserved KSPKKA motif indicated in a red box. (**B**) Protein extracts from BY-2 cell suspension culture were analysed by western blotting with the anti-H2A.W.6p antibody. Colorimetric and chemiluminescence images were overlaid using the ChemiDoc software to align the position of the signal with the protein ladder. (**C**) Flow cytometry profiles of tobacco BY-2 cell after aphidicolin block and release. Cells were analysed in a time course as in Figure 3A. BY-2 suspension culture was blocked with 20 µg/ml aphidicolin for 24 hours. Upon removal of the aphidicolin block, cells proceeded through the cell cycle in a synchronized manner that was followed by flow cytometry during a twelve hour-time course. (**D**) Control BY-2 cell suspension culture treated with 1% DMSO for 24 hours and then followed over a twelve-hour time course by flow cytometry. Only profiles for time points T0, T9, and T12 are shown. (**E**) Protein extracts from DMSO treated BY-2 cells were analyzed by western blotting with the indicated antibodies. Note that there is no fluctuation of phosphorylated H2A.W.6 (top panel) in unsynchronized cells compared to those synchronized by aphidicolin (see Figure 3A).

**S4 Fig.**
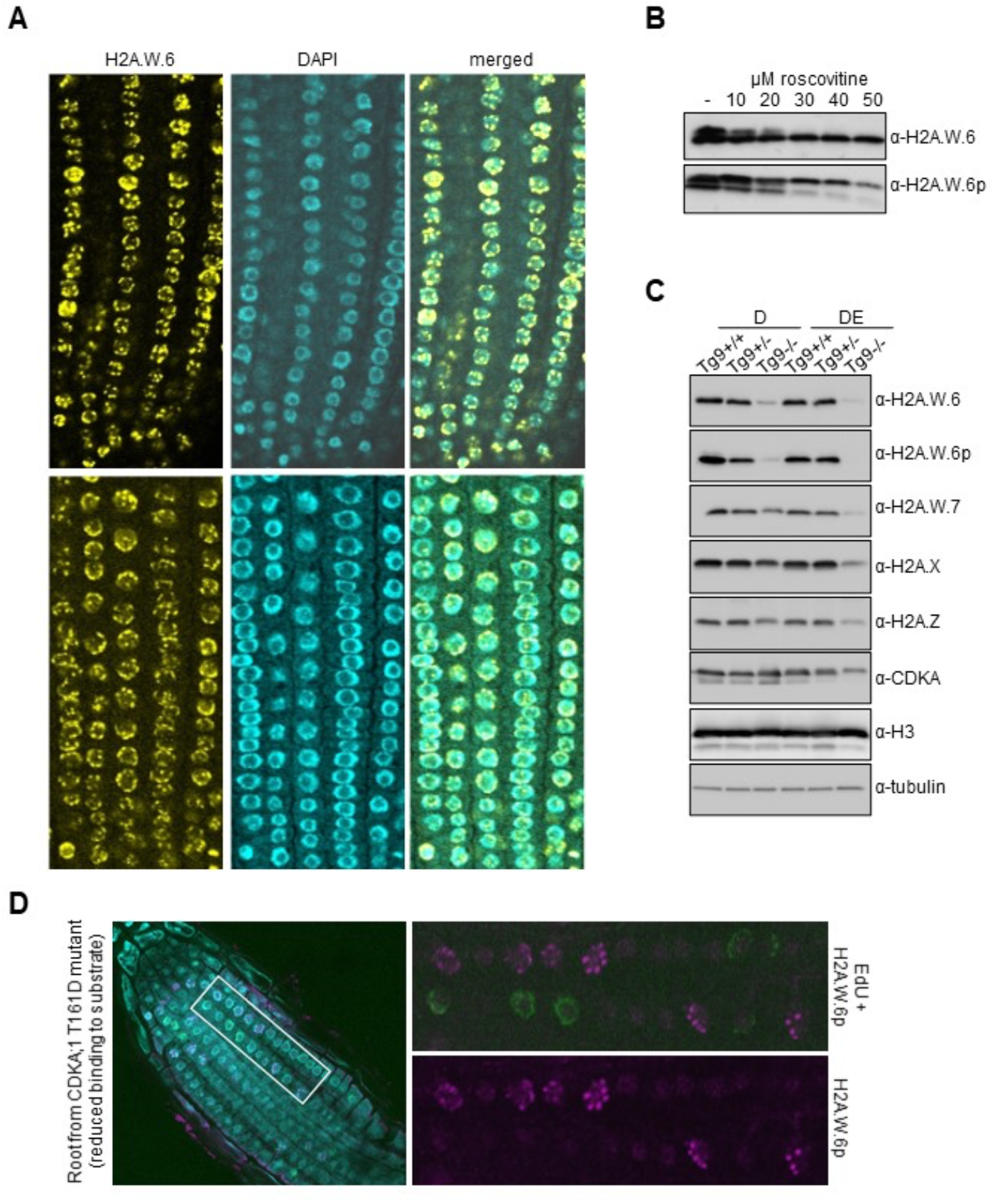
Phosphorylation of the H2A.W.6 KSPKK motif and expression of H2A.W.6 are linked to CDK activity in *Arabidopsis*. (**A**) Root tips of WT plants were immunostained with the H2A.W.6 antibody. Note that all nuclei display H2A.W.6 signal. Single confocal sections of two root tips are shown. (**B**) One-week old *Arabidopsis* cell suspension culture was treated for 24 hours with the indicated concentrations of roscovitine, a potent inhibitor of cell cycle-dependent kinases. Protein extracts were analyzed by western blotting with H2A.W.6 and H2A.W.6p specific antibodies. (**C**) Transgenic plants expressing weak hypomorphic mutants of CDKA;1, named D and DE, in the *cdka;1* mutant background [51, 52], were genotyped to identify heterozygous and homozygous plants (Tg9+/- and Tg9-/-). Nuclear protein extracts from 2-weeks old seedlings were analyzed by western blotting with the indicated antibodies. (**D**) CDKB;1, which is active at the G2/M transition, also phosphorylates the KSPKK motif of H2A.W.6. Root tips of plants expressing the CDKA;1 T161A hypomorphic mutant that results in reduced substrate binding were immunostained with the H2A.W.6p antibody (magenta) after EdU incorporation (green). Enlarged images on the right demonstrate the presence of H2A.W.6p in G2 nuclei but not in S phase nuclei labeled with EdU. A single confocal section is shown.

**S5 Fig.**
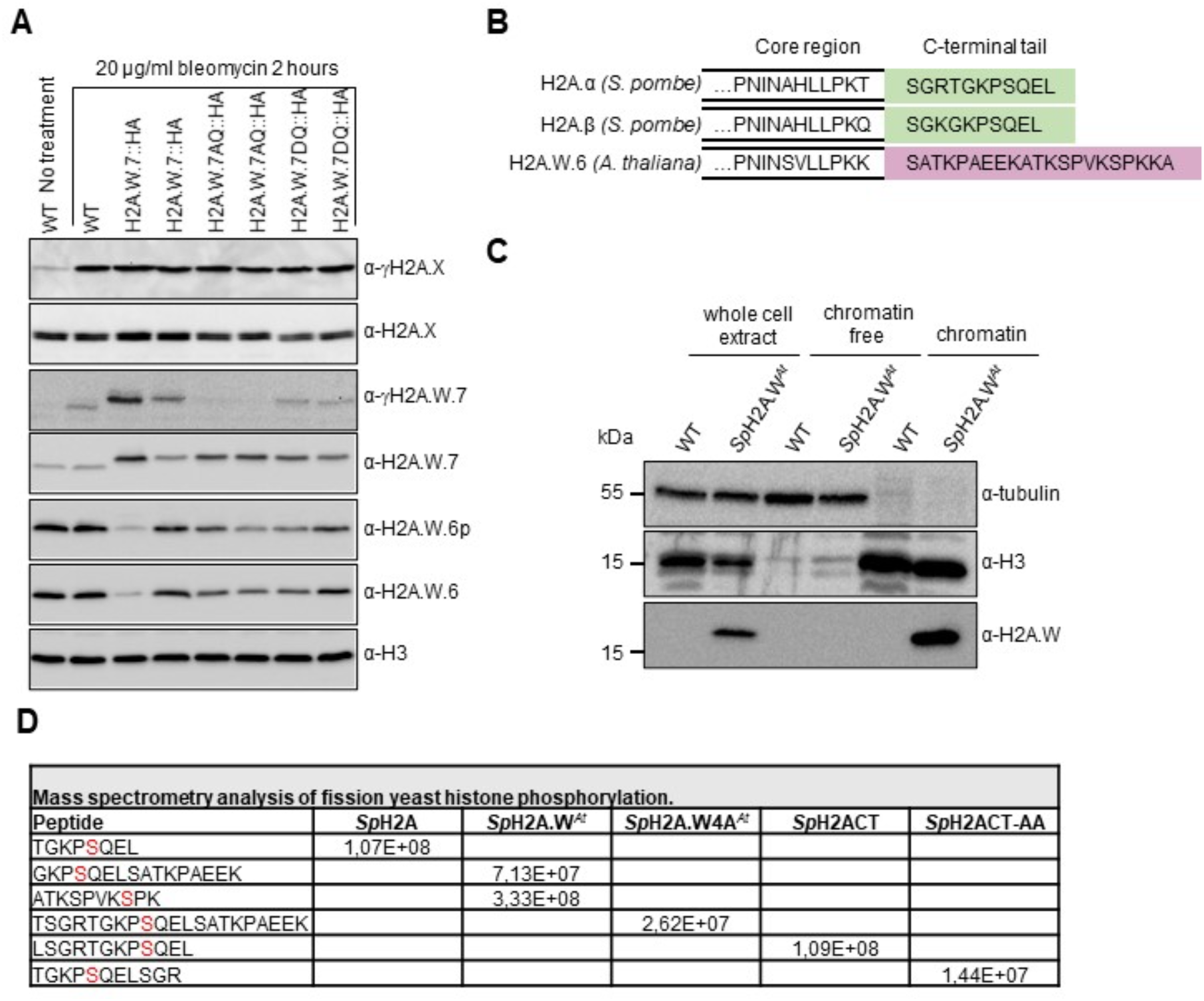
Analysis of KSPKK phosphorylation *in planta* and in *S. pombe*. (**A**) Analysis of γH2A.X, γH2A.W.7, and H2A.W.6p in transgenic seedlings expressing H2A.W.7 hypomorphic mutants in the SQ motif [16] after treatment with 20 µg/ml of bleomycin for 2 hours. Note that bleomycin treatment does not induce phosphorylation of the KSPKK motif on either WT H2A.W.7 or the AQ and DQ mutants of H2A.W.7; this band would be shifted above endogenous H2A.W.6p due to the presence of the HA tag. (**B**) Schematic diagrams of two *S. pombe* H2A variants and *Arabidopsis* H2A.W.6 with the indicated C-terminal tail sequences used for creation of the mosaic H2A variants depicted in Figure 4A. (**C**) Analysis of *Sp*H2A.W^*At*^ association with the chromatin. Whole cell, chromatin free, and chromatin bound fractions from *Sp*H2A and *Sp*H2A.W^*At*^ strains were analyzed by western blotting with the indicated antibodies. Tubulin and histone H3 were used as cytoplasmic and nuclear controls, respectively. (**D**) MS analysis of *Sp*H2A SQ motif phosphorylation in fission yeast strains expressing the indicated mosaic or mutated H2A variants. Highly similar levels, quantified as peak areas, of peptides covering phosphorylated SQ motif were measured in all strains.

**S6 Fig.**
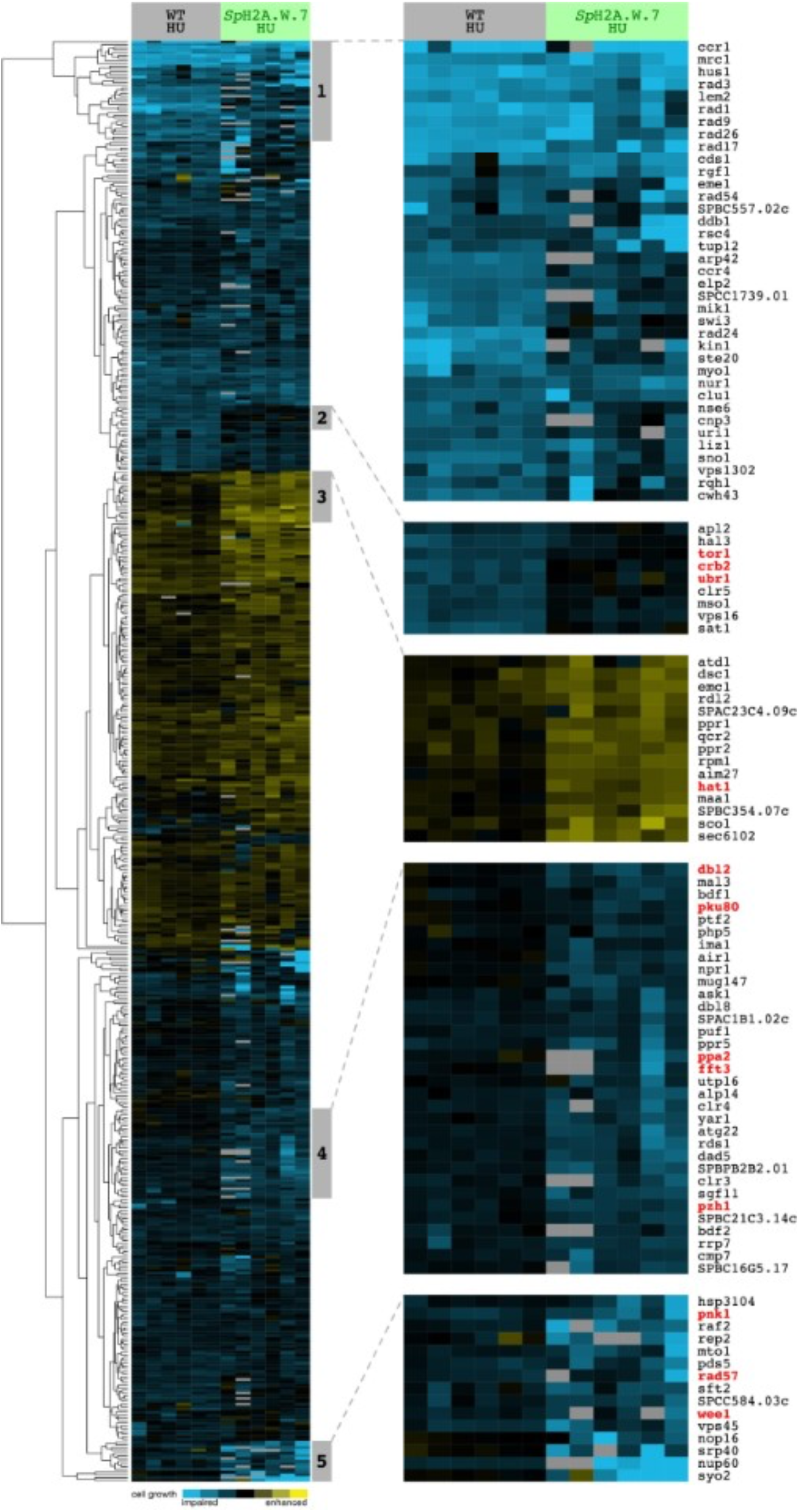
Cluster analysis of the SGA screen for deletion mutant that suppress growth of spH2A.WAt in 8mM HU plates.

